# Genome-Wide Profiling of tRNA Using an Unexplored Reverse Transcriptase with High Processivity

**DOI:** 10.1101/2023.12.09.569604

**Authors:** Yuko Nakano, Howard Gamper, Henri McGuigan, Sunita Maharjan, Zhiyi Sun, Keerthana Krishnan, Erbay Yigit, Nan-Sheng Li, Joseph A. Piccirilli, Ralph Kleiner, Nicole Nichols, Ya-Ming Hou

## Abstract

Monitoring the dynamic changes of cellular tRNA pools is challenging, due to the extensive post-transcriptional modifications of individual species. The most critical component in tRNAseq is a processive reverse transcriptase (RT) that can read through each modification with high efficiency. Here we show that the recently developed group-II intron RT Induro has the processivity and efficiency necessary to profile tRNA dynamics. Using our Induro-tRNAseq, simpler and more comprehensive than the best methods to date, we show that Induro progressively increases readthrough of tRNA over time and that the mechanism of increase is selective removal of RT stops, without altering the misincorporation frequency. We provide a parallel dataset of the misincorporation profile of Induro relative to the related TGIRT RT to facilitate the prediction of non-annotated modifications. We report an unexpected modification profile among human proline isoacceptors, absent from mouse and lower eukaryotes, that indicates new biology of decoding proline codons.

## Introduction

Transfer RNAs (tRNAs) comprise 70-90 nucleotides that fold into a stable L-shaped tertiary structure that embodies the genetic code. In this embodiment, an anticodon triplet is placed at the one end of the L while an amino acid is attached to the other end, providing a physical link that translates the corresponding codon into the amino acid building block during protein synthesis on the ribosome. To ensure the fidelity of protein synthesis, the L-shaped structure of each tRNA in higher eukaryotes is post-transcriptionally modified at 10-20% of the nucleobases^1^. These modifications are structurally and functionally diverse, with those in the anticodon loop modulating the quality of tRNA anticodon-codon pairing, while those elsewhere regulating tRNA stability, folding, and cellular localization^2, 3^. Importantly, because protein synthesis is tightly coupled to cell fitness^4–7^, both the abundance and the status of modifications of each tRNA respond to environmental cues^8, 9^. Indeed, expression levels of tRNAs, including those of the same amino-acid identity but different anticodon (i.e., isoacceptors) and those of the same anticodon but different sequence (i.e., isodecoders), are differentially regulated in tissues and in cell types^10–14^. Additionally, while some modifications in tRNAs are static, others are dynamic, changing levels in stress, across cell cycle stages, and in disease^3, 15–20^. Some modifications are even reversible, subject to the systems-level control of the writer and eraser enzymes that regulate their synthesis and removal^16^. The complexity of tRNA biology is further expanded by processing into 3’- and 5’-tRNA fragments^21–25^, which regulate RNA silencing and epigenetic expression^25–27^.

To evaluate the changes of cellular tRNA pools, multiple sequencing approaches have been developed. Most of these utilize the Illumina platform^14, 28–37^, where a tRNA pool is reverse transcribed by an RT into a cDNA library, which is PCR amplified and sequenced. The detection of modifications, if present in the W-C face, is by the occurrence of premature RT stops or RT misincorporation^38^. A few approaches utilize the Nanopore platform^39, 40^, in which a tRNA pool is directly passed through nanopores and the sequence of each tRNA is read as successive ion-current signals. The detection of modifications is by changes of an ion current event^41–44^. Between the two, the Illumina platform has a much larger database accumulated over multiple tRNAseq data, providing information for almost all tRNA modifications that are “RT-readable”, which are those that occur in the W-C face and thus would affect the RT readout. The high throughput and high capacity of the Illumina platform, giving rapid information on genome-wide tRNA dynamics, is also non-paralleled compared to the nanopore platform or the emerging platform of sequencing based on liquid chromatography-mass-spectrometry (LC-MS)^45, 46^. Importantly, the central component of the Illumina platform is a processive and efficient RT that is required to prepare the cDNA library. The traditional RTs are highlighted by SuperScript III and SuperScript IV, which are recombinant variants of the native RT of Moloney Murine Leukemia virus (MMLV) with improved thermostability and processivity^38^. A newer generation of RTs are encoded by group-II introns, which are mobile ribozymes that self-splice from precursor RNAs to invade into new genomic DNA. In this retrotransposition cycle, the intron-encoded RTs must efficiently copy the intron-released ribozymes, which are large and highly structured, requiring the evolution of these RTs with high processivity, high fidelity, and a strong strand displacement activity^47^. As such, group-II intron RTs generally have better performances than viral RTs and provide attractive resources for new discoveries^48^. However, only two group-II intron RTs are in use for tRNA research – TGIRT^49, 50^ in mim-tRNAseq^34^ and Marathon^51, 52^ in tRNA structure-seq^36^ and in ALL-tRNAseq^35^, all in the Illumina platform.

The most important feature of the RT for a successful tRNAseq is to read through as many sequences as possible. The failure to read through creates a truncated cDNA that lacks the sequence information downstream and complicates quantitative assessment of the tRNA. The critical challenge, however, is to overcome the extensive modifications in the stable structure of tRNA, which have hindered the field from moving forward. Notably, each RT has an intrinsic “signature” at each modification – to stop, to misincorporate, or to jump over. This signature can vary depending on the chemical nature of the modification and its structural context. For example, while *N*^1^-methylation of guanosine (m^1^G) interferes with W-C pairing, it can occur at position 9, where the tRNA acceptor stem turns from the D stem to stack on the T stem, or at position 37, on the 3’-side of the anticodon triplet, or at both. Similarly, while the m^1^A methylation interferes with W-C pairing, it can occur at positions 9/14/16/58 of tRNA, where each has a distinct local structural environment that may confer a distinct RT readout. Thus, the mechanism that determines the signature of the RT at each readable modification is important, as well as how this mechanism would change with the reaction time and temperature, and how the misincorporation profile of the RT is determined. However, despite the importance and timeliness of these questions, the signature profile of the RTs in current protocols at each readable modification has remained poorly defined. Addressing this unmet need will permit a more reliable prediction and assessment of non-annotated modifications.

Here we address this unmet need by establishing Induro-tRNAseq in the Illumina platform, using the recently developed and commercially available Induro (NEB M0681) as the RT. Induro is a new member of group-II intron RTs and has shown higher processivity than SuperScript IV and TGIRT^53^. We focus on how Induro recognizes and reads through the extensive modifications in tRNA. Among current tRNAseq methods, only the TGIRT-based mim-tRNAseq has the complete modification profile^34, 35^, providing an unbiased benchmark to evaluate Induro-tRNAseq. We show that Induro-tRNAseq is simpler and more comprehensive than mim-tRNAseq^34^, generating high quality data in a minimal number of steps without the need to purify tRNA by gels (as in mim-tRNAseq^34^) or by beads (as in MSR-seq^37^). We then use Induro-tRNAseq to elucidate the readthrough mechanism of the RT at each readable modification, showing that it is both temperature- and time-dependent, and that the driver that maximizes readthrough is selective removal of RT stops, without altering the misincorporation rate. This mechanistic insight, not yet available for other RTs, enables users to optimize their reactions. We also produce the misincorporation profile of Induro at each readable modification and compare it to that of TGIRT^34^, providing two datasets for the same modification to strengthen the power of prediction. Additionally, we generate the first comparison of tRNA modification profiles between human and mouse, offering the potential to use mouse genetics to understand human diseases associated with mutations in tRNA modification enzymes. Based on our comparison between Induro and TGIRT, and our analysis of human and mouse profiles of tRNA modifications, we uncover an unexpected modification profile of human proline isoacceptors that is separately confirmed by LC-MS analysis. This modification profile is absent from mouse and from lower eukaryotes, suggesting that it entails a new mechanism of decoding proline codons in human biology.

## Results

### The multiplexing Induro-tRNAseq workflow

We developed Induro-tRNAseq with the following considerations. First, to reduce the number of experimental steps, we used total RNA as the input, rather than gel- or beads-purified tRNA. We isolated total RNA from cultured human cells and, for the focus of determining the modification profile, we removed their aminoacyl groups by a brief alkaline hydrolysis (Fig. 1a, step 1). We took advantage of the 3’-end CCA sequence of all mature tRNAs and optimized a splint ligation to attach a barcoded 3’-adaptor to the CCA end using T4 Rnl2 (Supplementary Table 1) (step 2). The ligation efficiency was high, typically reaching >75% and generating ligation products in the predicted size of ∼120 nucleotides (Fig. 1b). Several pools of tRNA, each with a specific barcode, were combined at this step to start a multiplexing workflow. We then used Induro to extend an RT primer (step 3) from a region common to barcodes of the multiplexed pools, generating a range of cDNA products (Fig. 1c), consistent with the presence of modifications that inhibited the RT from end-to-end cDNA synthesis. Second, we isolated all cDNA products in the range of 90-180 nucleotides, using gel purification (step 4), which was necessary to remove primer molecules that were extended without tRNA (Fig. 1c). These primers can be a major contaminant downstream, due to their ability to participate in both circularization and PCR amplification^54^. Third, the gel-purified cDNAs, which contained annealing sites for both the universal and multiplex PCR primers, were circularized by CircLigase to generate circular templates for PCR amplification (step 5). This circularization reaction is more efficient than ligating an adaptor to the 3’-end of the cDNAs. We next used the high-fidelity Q5 DNA polymerase in PCR amplification to add Illumina adaptors to the cDNA libraries (Supplementary Table 2) (step 6). The extended cDNA libraries, after a minimal cycles of PCR amplification (Fig. 1d), were isolated from a native PAGE and sequenced on an Illumina NextSeq 500 platform.

**Figure 1.**
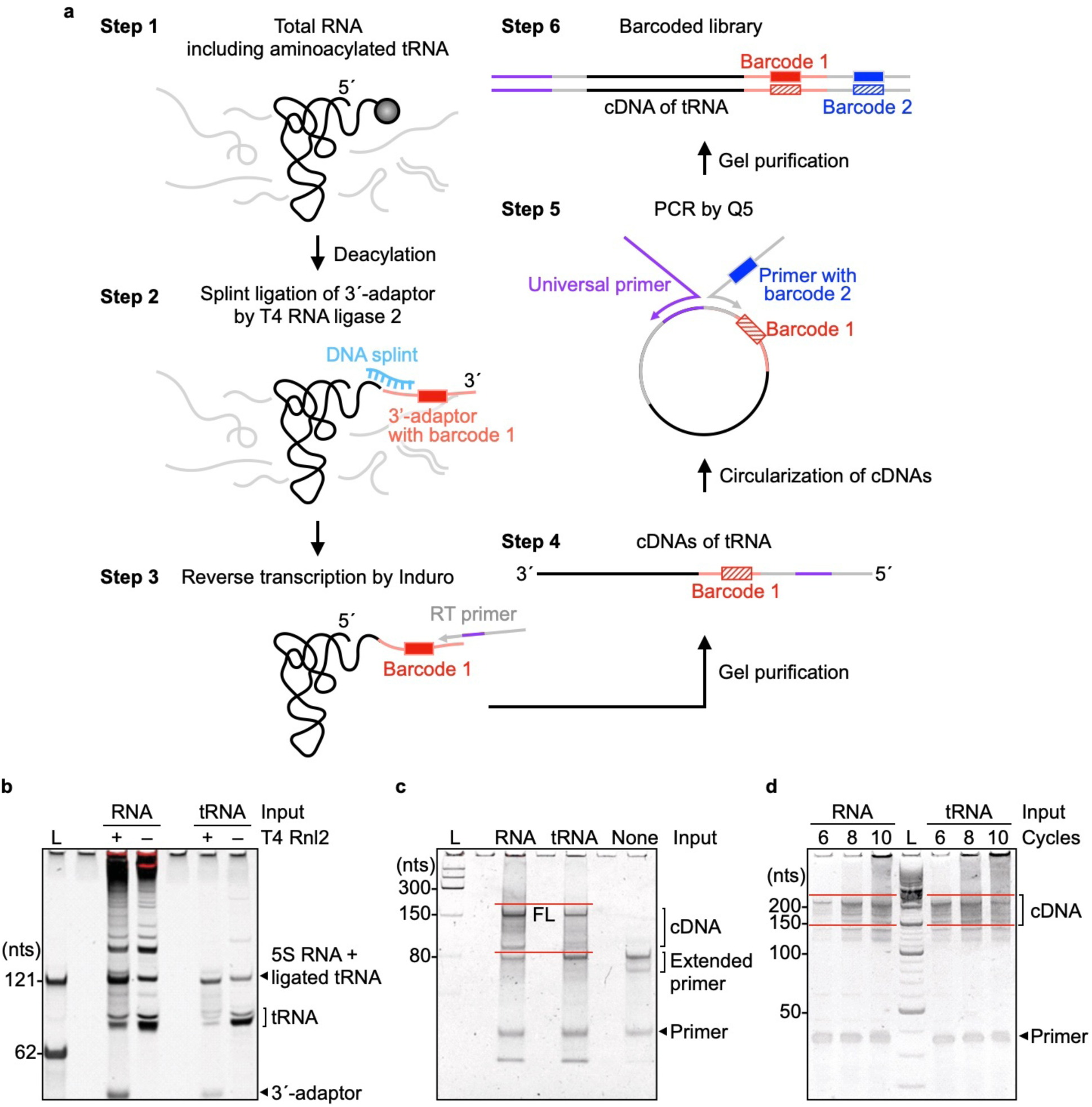
The Induro-tRNAseq workflow. **a.** Diagram of tRNA-seq library preparation. Step 1: Total RNA including charged aminoacyl-tRNAs, where the amino acid is depicted as a black ball, is extracted from mammalian cultured cells, or tissue samples, and used without purification of the tRNA pool unless otherwise specified. Charged tRNAs are deacylated in a glycine buffer, pH 9.0. Step 2: A 3’-barcoded adaptor is hybridized to a bridging DNA splint and is ligated to the 3’-end of deacylated tRNAs by T4 RNA ligase 2 (T4 Rnl2). Several pools of tRNA libraries can be combined here to start a multiplex workflow. Step 3: An RT primer is hybridized to the 3’-adaptor to initiate cDNA synthesis by Induro. Step 4: cDNA products are gel purified and circularized by Circligase. Step 5: A new primer with barcode 2 and a universal primer are used to PCR amplify circularized cDNAs by Q5 DNA polymerase. Step 6: The final barcoded library contains tRNA sequence (black), barcode 1 (red), and barcode 2 (blue). **b.** A denaturing gel image of tRNA ligated with the 3’-adaptor after step 2. Total RNA isolated from human cultured cells or a gel-purified tRNA pool was used as the input material. L: 62-mer and 121-mer DNA markers. **c.** A denaturing gel image of cDNA products after reverse transcription by Induro in step 3. cDNA products of 90 - 180 nts were excised and extracted from the gel. L: small range RNA ladder (NEB); none: a control RT reaction without an input RNA; FL: cDNA of full-length tRNA. **d.** A non-denaturing gel image of the cDNA library after PCR amplification in step 5. cDNAs of 150 - 240 base pairs (bps) were excised and extracted. L: O’RangeRuler 10 bp DNA ladder + GeneRuler 50 bp DNA ladder (Thermo Fisher). All gels were stained with SYBR gold.

A side-by-side comparison of the workflow, using total RNA vs. purified tRNA pool as input, showed similar performance throughout. While the tRNA pool might have generated a better yield of splint ligation (Fig. 1b), the total RNA sample led to a better yield of the Induro-generated full-length cDNA product (Fig. 1c), although these differences were minor. Notably, in the final step of generating PCR-amplified cDNA libraries for sequencing, the two samples had virtually identical yields (Fig. 1d). Thus, using total RNA isolated from cells, rather than purified tRNA, provides a simpler method to start the workflow. Additionally, if the focus were to determine the level of charged tRNA, the protocol can be modified to keep the aminoacyl groups and to subject the pool to a periodate reaction to remove uncharged tRNA^37^.

We isolated total RNA from cultured human K562 and HEK293T cells to compare the performance between Induro-tRNAseq and mim-tRNAseq, using data from the latter for both cell lines^34^. The barcoded libraries from both cell lines generated 2-7 million total raw reads, which were trimmed to remove the adaptor and the 5’-RN sequences (R: purine, N: a mixture of all four natural nucleotides) that were introduced during the RT reaction^34^. The processed reads were aligned to the high confidence set of human tRNA sequences annotated in gtRNAdb of the hg38 reference genome^55^, allowing mismatches up to 10% of each sequence. To detect non-annotated modifications, mapping was repeated with an updated SNP index allowing new mismatches derived from the first round of alignment. To recover reads that terminated at m^1^A58 in the T loop, the minimum score was set at a threshold that allowed mapping of reads of at least 10 nucleotides. From both cell lines, ∼90% of total reads were uniquely mapped to gtRNAdb sequences, closely similar to that in mim-tRNAseq^34^ and similar between samples starting with total RNA or with purified tRNA pool (Supplementary Fig. 1a). This high frequency of uniquely mapped reads demonstrates the high quality of reads and the lack of contaminating sequences or inappropriate alignment. It also validates the robustness of the workflow and the utility of starting with total RNA. Histograms of mapping quality (MAPQ) scores using total RNA as the input showed that all reads were in the range of 36-40 (Supplementary Fig. 1b), indicating high mapping confidence. Reproducibility between biological replicates was excellent, with the coefficient of determination *r*^2^ for each replicate = 0.9999.

### Profiling Induro for readthrough of modifications

We profiled Induro for readthrough of RT-readable modifications. In the canonical tRNA sequence framework, these modifications are m^1^A9/58, m^1^G9/37, acp^3^U20 (acp^3^U = 3-amino-3-carboxy propyl-uridine at position 20), m^2^ G26 (m^2^ G = *N*^2^, *N*^2^ -dimethyl guanosine at position 26), m^3^C32 (m^3^C = *N*^3^-methyl cytidine at position 32), I34 (I = inosine at position 34), m^1^I37, yW37 (yW = wybutosine at position 37), ms^2^i^6^A37 and ms^2^t^6^A37 (ms^2^i^6^A = 2-methylthio-*N*^6^-isopentenyl-adenosine at position 37; ms^2^t^6^A = 2-methylthio-*N*^6^-threonyl-carbamoyl-adenosine at position 37) (Fig. 2a). For each isodecoder tRNA, we calculated the readthrough frequency by the read counts of both correct and incorrect nucleotide incorporation over the total counts, including RT stops. While modifications that occur outside of the W-C face (e.g., ψ (pseudouridine), D (dihydrouridine), m^5^C, 1m^5^U (5-taurine-methyl-uridine), and T (ribo-thymine)) are currently RT-unreadable, they may become readable after chemical treatments, although we did not consider them in this study, due to the potential bias of chemical reactions.

**Figure 2.**
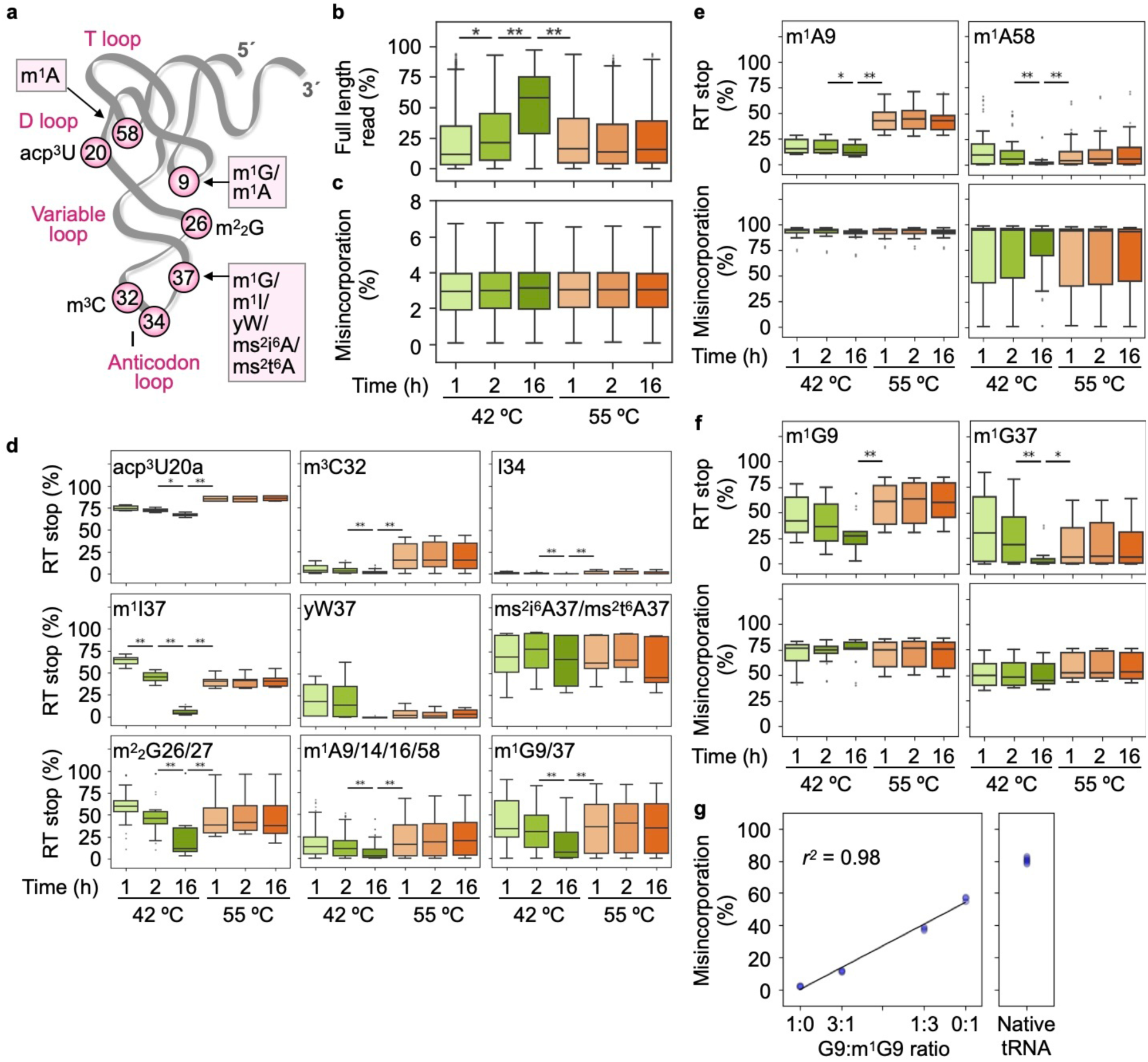
Profiling Induro reactions. **a.** The L-shaped structure of tRNA showing RT-readable modifications. **b.** Boxplots of the yield of full-length cDNA (%) indicating end-to-end cDNA synthesis at 42 °C (green) or 55 °C (orange) after the RT reaction for 1, 2, or 16 h. Data were collected from total RNA of K562 cells as the input (n = 2; technical replicates). **c.** Boxplots of the frequency of misincorporation (%) in each RT reaction. **d.** Boxplots of the frequency (%) of RT stops in a 3-nucleotide window centering on each annotated modification at position 0. **e.** Boxplots of the frequency (%) of RT stops and RT misincorporation at m^1^A9 and m^1^A58. **f**. Boxplots of the frequency (%) of RT stops and RT misincorporation at m^1^G9 and m^1^G37. Center line: median; box limits: upper and lower quartiles; whiskers: 1.5X interquartile range; points: outliers. **g.** Left: A scatter plot showing the frequency (%) of RT misincorporation as a function of the G9:m^1^G9 ratio of the transcript of human mt-Leu(TAA) (n = 2; technical replicates). Right: The frequency of misincorporation of the native mt-Leu(TAA) in sequencing analysis of a total RNA sample. Data were collected from K562 cells with total RNA as the input under the RT reaction condition at 42 °C for 16 h (n = 2; technical replicates). The Pearson correlation coefficient is indicated by *r*^2^. Student’s *t*-test was performed by a two-sided analysis (\**p* < 0.1, \*\**p* < 0.01).

We began by testing both the temperature and time of the Induro reaction to identify the condition that maximized its readthrough. While the recommended temperature of Induro is 55 °C (NEB M0681), we also tested 42 °C to explore the possibility of lowering the enzyme dynamics at each modification to promote readthrough. Additionally, while Induro is capable of readthrough of a 12 kb RNA in less than 10 min at 55 °C (NEB M0681), we tested its readthrough of tRNA by monitoring at 1, 2, and 16 h. Unexpectedly, while the frequency of readthrough to produce the full-length yield was low (10%) at 55 °C, it did not improve over time, whereas the frequency increased to 60% at 42 °C over 16 h (Fig. 2b). Notably, an increase of readthrough is an interplay between the increase of misincorporation and the concomitant decrease of RT stops. Intriguingly, measurement of the overall misincorporation rate showed that it was consistently low (3%) across temperature and time (Fig. 2c), suggesting that the increase in readthrough is entirely driven by decreases of RT stops. The specific increase in readthrough at 42 °C, but not at 55 °C, supports the notion of reduced dynamics of the RT to suppress its falling-off from the tRNA template. The low frequency of misincorporation at the global level indicates that the RT is largely faithful, while the consistency of the frequency indicates that the RT maintains a relatively fixed ratio of correct vs. incorrect nucleotide incorporation regardless of the reaction condition. Importantly, despite the low frequency of readthrough at 55 °C, the uniquely mapped reads remained high (83-89%) (Supplementary Fig. 1c), supporting the high quality of reads even with lower number of reads.

We next determined the mechanism of increased readthrough by reduction of RT stops and whether the reduction was uniform at all modifications or was site-specific. We first developed a comprehensive map of the interplay between RT stops and RT misincorporation at each readable modification (Supplementary Fig. 2a). At the modification of interest, defined as position 0, we only found RT stops or RT misincorporation from –1 to +1, despite an exhaustive search from positions –2 to +2, indicating the lack of RT jump-over. We also confirmed that, relative to each position 0, there was no adjacent modification that would have caused RT stops, allowing precise assignment of RT stops. The map showed a diverse interplay between RT stops and RT misincorporation, indicating the idiosyncrasy of Induro in response to different modifications. In most cases, Induro responded primarily by misincorporation at position 0, although it also responded by a low level of RT stops (at m^1^A9/14/16/58, m^1^G9/37, m^2^_2_G26/27, m^3^C32, I34, m^1^I37, and yW37). Only in two cases did the enzyme respond exclusively by RT stops (at acp^3^U20 and ms^2^i^6^A37/ms^2^t^6^A37). The predominant representation of RT misincorporation relative to RT stops indicates an overall high propensity of readthrough. Notably, we observed site-to-site variation of both RT misincorporation (a near 100% frequency at m^2^_2_G26/27 and I34) and RT stops (a spread-out frequency at –1, 0, and +1 at acp^3^U20, but solely at position +1 at ms^2^i^6^A37/ms^2^t^6^A37). Using this comprehensive map, we then showed that RT stops indeed decreased over time at all modifications where stops were detected, and that this decrease was specific at 42 °C (Fig. 2d). In contrast, the frequency of misincorporation remained the same at all sites across temperature and time (Supplementary Fig. 2b). Combined, these results clearly show that it is the decrease of RT stops, with site-specific degree, that drives the increase of readthrough over time (Fig. 2b).

We developed a higher resolution map for m^1^A and m^1^G and found a position-dependent decrease of RT stops. While other modifications also occur at multiple sites, their frequency was much lower (e.g., m^2^ G26/27). For both m^1^A and m^1^G, the decrease of RT stops with time was milder at position 9 relative to position 58 and 37, respectively (Fig. 2e, f). This positional effect can be explained by the tRNA L-shaped structure, where position 9 is in a tight turn with multiple tertiary interactions that stabilize the turn, whereas positions 37 and 58 are both in a loop in a less structured environment (Fig. 2a). The differential reduction of RT stops at position 9 relative to positions 37 and 58 suggests that the enzyme responds to structural barriers of each modification. Notably, the degree of decrease of RT stops, while specific to the position and chemical structure of each modification, was only observed at 42 °C over time, consistent with the temperature-specific effect (Fig. 2b).

The time-dependent increase in readthrough specifically at 42 °C was observed not only for cytosolic tRNAs (cyto-tRNAs), but also for mitochondrial tRNAs (mt-tRNAs) (Supplementary Fig. 2c, d). Also, while readthrough increased, the misincorporation rate remained constant for both classes of tRNAs (Supplementary Fig. 2c, d). These results strengthen the notion that the time-dependent increase in readthrough at 42 °C is entirely driven by reduction of RT stops.

To quantify the stoichiometry of each modification, we noted the near 100% frequency of RT misincorporation at m^2^_2_G26/27 and I34 (Supplementary Fig. 2b), indicating that each is homogeneously modified. Thus, a fractional reduction of RT misincorporation at each site would represent a fractional loss of the modification, which can occur in a specific cellular condition. For other modifications, the stoichiometry cannot be determined solely by the frequency of RT stops or RT misincorporation but can be determined by a calibration analysis. Using the m^1^G9 methylation in the native mt-Leu(TAA)^56^ as an example, where TAA is the gene for the anticodon, we prepared a transcript of the tRNA, lacking any modification, and a separate transcript containing m^1^G9 as the single modification at 100%. Sequencing analysis of the two transcripts, mixed at different ratios, revealed a linear correlation between the frequency of misincorporation and the stoichiometry of m^1^G9, with the Pearson *r*^2^ of 0.98 (Fig. 2g), showing the quantitative precision of the workflow. Interestingly, we found that the RT misincorporation frequency at saturating m^1^G9 reached 60% with the transcript tRNA but reached 80% with the native tRNA containing the full complement of all natural modifications (Fig. 2g), validating the notion that the RT is sensitive to the local environment of each modification. Indeed, in contrast to the transcript tRNA, the native tRNA contains m^2^G10 following m^1^G9 (ref^56^), suggesting that the adjacent modification has increased the RT misincorporation frequency at m^1^G9.

Combined, our profiling of Induro demonstrates its ability to read through all but two of the RT-readable modifications in tRNA. Maximum processivity of readthrough occurred at 42 °C overnight in a mechanism that removes site-specific RT stops without altering the RT misincorporation rate. All subsequent reactions were performed in the condition that maximized readthrough.

### Comprehensive coverage of tRNA by the Induro-tRNAseq workflow

While the workflow starting with total RNA or with purified tRNA (as in mim-tRNAseq^34^) performed similarly overall (Fig. 1d), we investigated whether the former would have additional advantage besides supporting a simplified procedure. We began with analysis of the conserved CCA triplet at positions 74-76, where the terminal A76 is the universal site for aminoacylation. While the CCA triplet does not affect the modification profile of each sequence, it is protruded from the tRNA structure and is most sensitive to degradation, suggesting that its integrity is a measure of the tRNA 3’-end quality^57^. Analysis of the fraction of incomplete CCA in each isoacceptor family, including 3’-N, 3’-NC, and 3’-NCC, where N is the discriminator base at position 73, relative to all 3’-end sequences, showed similar quality either starting with total RNA or with purified tRNA. This is shown in samples isolated from K562 (6.2% for total RNA and 4.1% for the tRNA pool) (top two rows, Fig. 3a), and in samples isolated from HEK293T cells (6.8% for total RNA and 7.5% for the tRNA pool) (bottom two rows). Thus, the two starting methods to begin the workflow had no major difference in producing high-quality 3’-ends.

**Figure 3.**
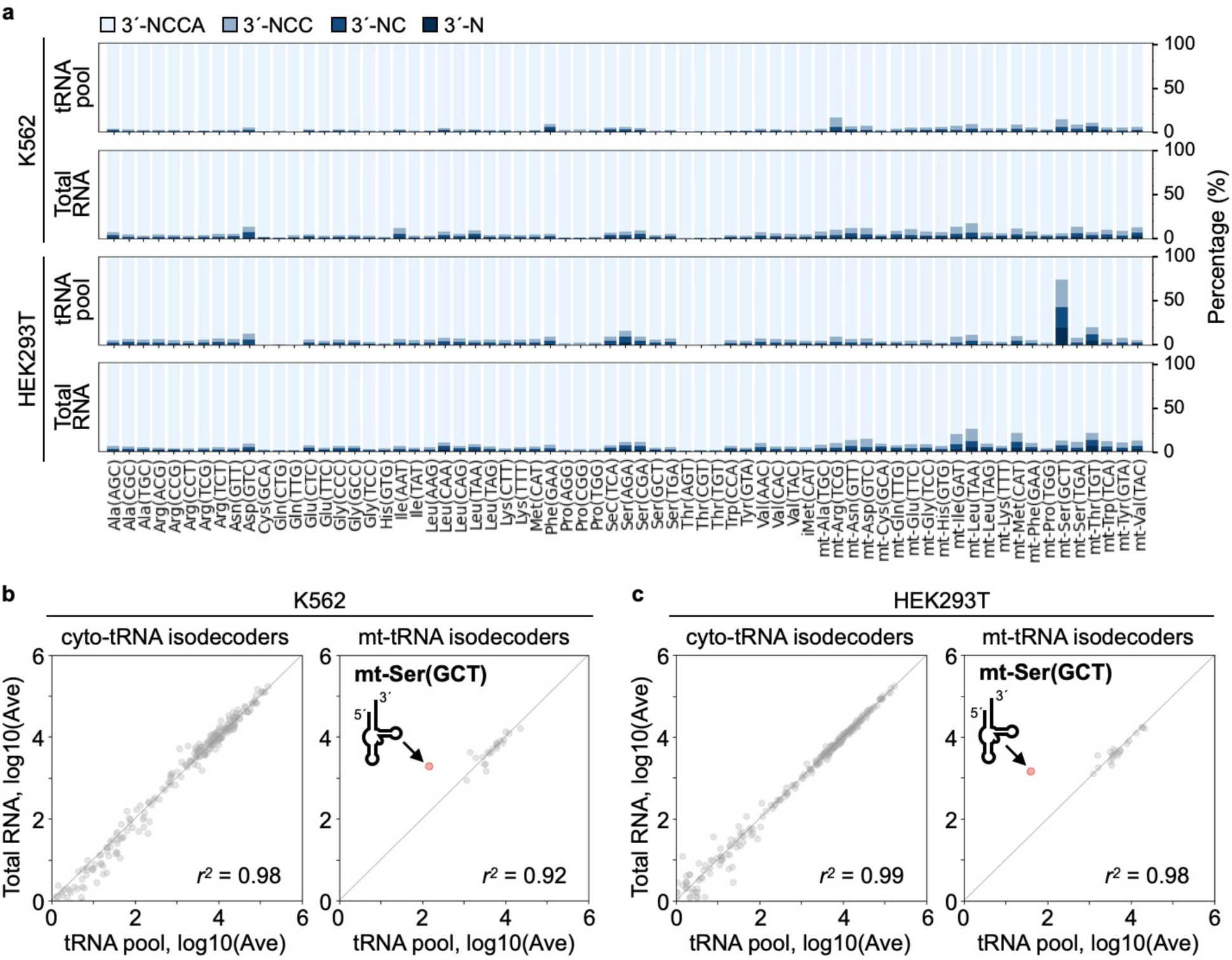
The workflow starting with total RNA or with a purified tRNA pool. **a.** Analysis of the 3’-CCA end, showing cumulative bar graphs of the average % of each state 3’-NCCA, 3’-NCC, 3’-NC, and 3’-N with increasing density of the blue color. Total RNA (n = 4; biological replicates) and a purified tRNA pool (n = 2; biological replicates) were analyzed for K562 cells, and total RNA (n = 2, biological replicates) and a tRNA pool (n = 2; biological replicates) were analyzed for HEK293T cells. **b.** Scatter plots indicate differential abundance of cyto-tRNA and mt-tRNA isodecoders of K562 cells. Total cyto-tRNA isodecoders (279, 276 in two duplicates) and mt-tRNA isodecoders (22 in each sample) were analyzed. **c.** Scatter plots demonstrate differential abundance of cyto-tRNA and mt-tRNA isodecoders of HEK293T cells. Total cyto-tRNA isodecoders (273, 275) and mt-tRNA isodecoders (22 in each sample) were analyzed. Shown in each comparison are log-transformed read counts normalized by DESeq2 with the Pearson correlation coefficient indicated by *r*^2^. The cloverleaf structure of mt-Ser(GCT) pointing to the higher abundance in the total RNA sample relative to the purified tRNA pool is shown in **b**, **c**.

In contrast, the workflow starting with total RNA had a much more comprehensive coverage of all tRNA sequences. In samples of K562 cells, while the relative abundance of cyto-tRNAs was similar between the total RNA sample and the purified tRNA pool (Pearson *r*^2^ of 0.98), one specific tRNA, mt-Ser(GCT), was detected at 13-fold more abundant in the total RNA sample relative to the purified tRNA sample (Pearson *r*^2^ of 0.92) (Fig. 3b). Similarly, in samples of HEK293T cells, while the relative abundance of cyto-tRNAs was similar between the total RNA sample and the purified tRNA sample (Pearson *r*^2^ of 0.99), mt-Ser(GCT) was detected as 38-fold more abundant in the total RNA sample (Pearson *r*^2^ of 0.98) (Fig. 3c).

Notably, mt-Ser(GCT) is unique among all human tRNAs as having the shortest sequence (62 nucleotides), lacking the entire D stem-loop^56^ (Fig. 3b, c). We attribute the detection of higher abundance of this tRNA in the total RNA sample to the inclusiveness of the method of all tRNA species, regardless of the size. The short sequence of mt-Ser(GCT) would have been lost from the purified tRNA pool isolated by gel-size selection of 60-100 nucleotides. Thus, starting with total RNA has the important advantage in isolating tRNA species with a diverse size range. This can be useful for analysis of 3’-half tRNA fragments (20-35 nucleotides) or tRNA-aptamer fusions (up to 200 nucleotides)^58^, the latter of which can be engineered to image tRNA translocation across the cellular space.

### Similarities and differences between Induro and TGIRT

We used the datasets generated from Induro-tRNAseq and from mim-tRNAseq^34^ to elucidate the similarities and differences between Induro and TGIRT, respectively. The two methods, except for the differences in RT and the input material, are similar overall. This comparison is important to provide information on Induro vs. TGIRT as two members of the group-II intron family to evaluate their relatedness. We confirmed that the two sequencing methods showed similar abundance profiles of cyto-tRNAs based on similar numbers of detectable species (Pearson *r*^2^ of 0.95) (Fig. 4a). We also confirmed that the two sequencing methods showed similar abundance profiles of mt-tRNAs, except for mt-Ser(GCT), which was under-represented in mim-tRNAseq due to the use of purified tRNA as the input in the latter. Indeed, the more comprehensive coverage of our workflow relative to mim-tRNAseq is shown as a 423-fold more abundance of mt-Ser(GCT) in samples of K562 cells (Pearson *r*^2^ of 0.85 based on 22 mt-tRNAs) (Fig. 4b) and a 11-fold more abundance in samples of HEK293T cells (data not shown). This emphasizes the importance of starting the workflow with total RNA, rather than with purified tRNA.

**Figure 4.**
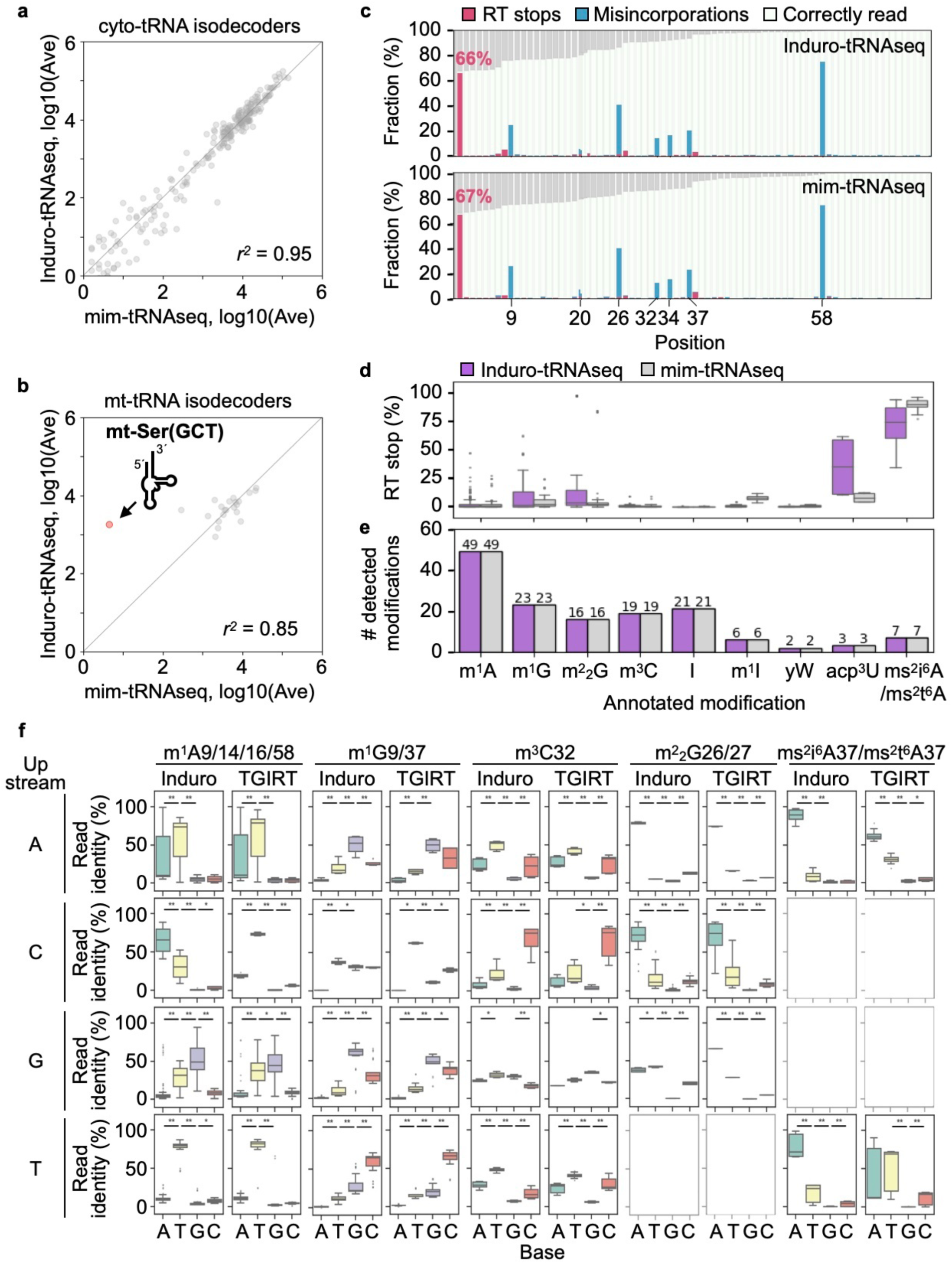
Induro-tRNAseq vs. mim-tRNAseq. **a.** A scatter plot indicates differential abundance of cyto-tRNA isodecoders of K562 cells by Induro-tRNAseq and by mim-tRNAseq. The number of cyto-tRNAs detected was 279 and 262, respectively. **b.** A scatter plot indicates differential abundance of mt-tRNA isodecoders of K562 cells by Induro-tRNAseq and by mim-tRNAseq. The number of mt-tRNAs detected was 22 in both workflows. Shown in each comparison are log-transformed read counts normalized by DESeq2 and the Pearson *r*^2^ in each. **c.** Cumulative bar graphs show the frequency (%) of RT stops (pink) and RT misincorporations (blue) at each readable modification as averaged from all tRNA isodecoders isolated from K562 cells, as well as the frequency (%) of correct reads (white) and of loss of coverage (gray). RT stops were determined by the number of stops at a given position relative to the total number of reads. **d.** Boxplots of the frequency (%) of RT stops at each readable modification as detected by Induro-tRNAseq and by mim-tRNAseq. Center line: median; box limits: upper and lower quartiles; whiskers: 1.5X interquartile range; points: outliers. **e.** A bar graph indicates the number of reads at each readable modification as detected from all tRNA species by Induro-tRNAseq and by mim-tRNAseq. **f.** Boxplots of the distribution of read identity (in %) of each readable modification as detected from tRNA of K562 cells by Induro-tRNAseq and by mim-tRNAseq in response to different upstream nucleotides. The analysis of Induro-tRNAseq was based on this work (n = 4, biological replicates), while that of mim-tRNAseq was extracted from the published dataset^34^. Center line: median; box limits: upper and lower quartiles; whiskers: 1.5X interquartile range; points: outliers. Student’s *t*-test was performed by a two-sided analysis (\**p* < 0.1, \*\**p* < 0.01).

The two sequencing methods were also similar in quantitative detection of RT-readable modifications in tRNA, showing essentially identical frequencies of misincorporation across the sequence framework (Fig. 4c). While this suggests that the two RTs have a similar intrinsic fidelity of reading each modification, some differences exist. The most notable was at the two modifications that cause exclusive RT stops for Induro – acp^3^U20 and ms^2^i^6^A37/ms^2^t^6^A37 (Supplementary Fig. 2a). While both RTs stalled at the two modifications, Induro stopped with a higher frequency relative to TGIRT at acp^3^U20, but with a similar frequency at ms^2^i^6^A37/ms^2^t^6^A37 (Fig. 4d). Despite this, the two RTs detected the same number of modifications in the respective workflows (Fig. 4e).

While Induro and TGIRT share a similar misincorporation frequency at each readable modification (Fig. 4c), we determined whether they differ in the misincorporation profile to produce a distinct “read identity” of each modification. We define the read identity as the direct sequencing read of the RT misincorporation in response to a modification. For example, a misincorporation of C in response to m^1^A would produce a read identity of G as the complement to C.

We found a subset of modifications whose read identity is context-independent, such as acp^3^U20a, I34, m^1^I34, and yW37 (Supplementary Fig. 3a). Of these, yW37 is the only one that elicited a difference between the two RTs, where Induro produced primarily G as the read identity whereas TGIRT produced primarily T. We also found another subset of modifications whose read identity is context-dependent, such as m^1^A9/14/16/58, m^1^G9/37, m^3^C32, m^2^_2_G26/27, and ms^2^i^6^A37/ms^2^t^6^A37. Of these, the two RTs showed more differences in response to the upstream than the downstream nucleotide (Fig. 4f, Supplementary Fig. 3b). We present data for each modification by combining reads at all positions where it occurred in the tRNA sequence, which would be dominated by those at the most prominent positions (e.g., m^1^A9 and m^1^A58 relative to m^1^A14 and m^1^A16). Thus, in response to the upstream nucleotide C of the combined m^1^A9/14/16/58 modifications, while Induro produced A as the read identity, TGIRT produced T (Fig. 4f). Two other examples of differences are also notable (Fig. 4f). In response to the upstream nucleotide G of the combined m^2^ G26/27 modifications, while Induro produced a mixture of A/T as the read identity, TGIRT produced A. In response to the upstream nucleotide U of the combined ms^2^i^6^A37/m^2^t^6^A37 modifications, while Induro produced A as the read identity, TGIRT produced

T. In contrast, only one clear difference existed in response to the downstream nucleotide. In response to the downstream nucleotide A of the combined m^2^ G26/27 modifications, while Induro produced a mixture of A/T as the read identity, TGIRT produced A (Supplementary Fig. 3b).

At m^1^A and m^1^G, while both Induro and TGIRT were sensitive to positional effects of each chemical structure, they however showed a similar response (Supplementary Fig. 3c). Both produced T as the read identity of m^1^A9 but a mixture of T/G for m^1^A58. Similarly, both produced C as the read identity of m^1^G9 but a mixture of G/C/T for m^1^G37. Combined, the similarities and differences between the two RTs constitute a wealth of new information that can be explored and developed to enhance read identity in profiling tRNA modifications.

### Induro-tRNAseq quantifies a disease-associated loss of tRNA

We tested the ability of Induro-tRNAseq to detect changes in tRNA profile in tissue samples. We used the mouse Arg(TCT)-4 as an example (Fig. 5a), which is a brain-specific cytosolic Arg isodecoder, where the C50U mutation in the T loop decreases pre-tRNA processing and consequently the abundance of the mature tRNA^59^. This C50U mutation is an underlying cause of neurodegeneration, due to ribosome stalling upon loss of the mature tRNA at the molecular level^59^. We performed Induro-tRNAseq using the brain tissue of a normal mouse (the B6N strain) and a mutant mouse (the B6J strain) harboring the C50U mutation. We readily detected the loss of Arg(TCT)-4 in the mutant brain among a similar number of cyto-isodecoders (Pearson *r*^2^ of 0.98), but similar abundances relative to the normal brain among 22 mt-isodecoders (Pearson *r*^2^ of 0.97) (Fig. 5b). The loss of Arg(TCT)-4 from the mutant brain was 25-fold, which was a more severe loss than the 10-fold loss reported previously by Northern blot analysis^59^.

**Figure 5.**
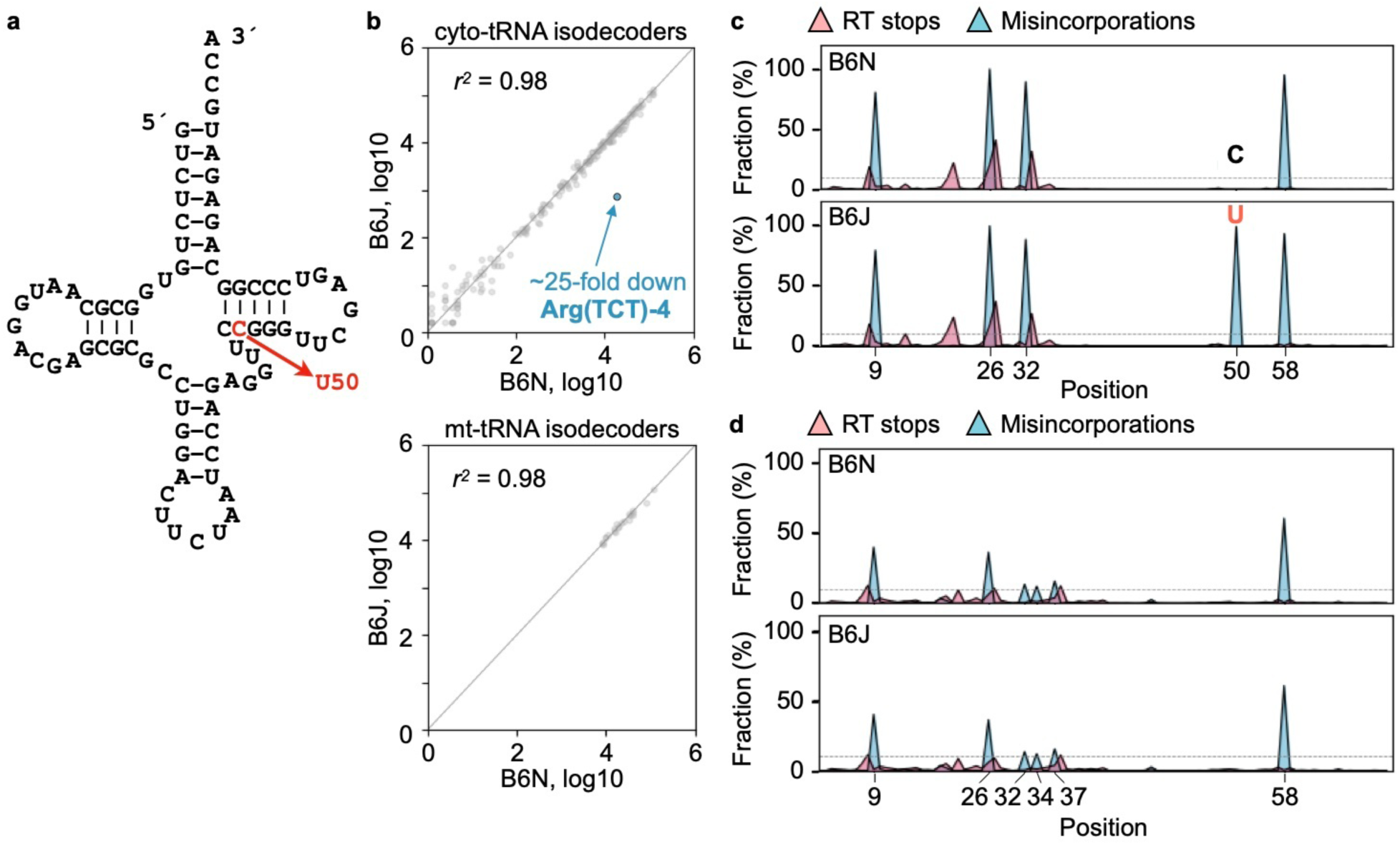
Induro-tRNAseq analysis of total tRNA from mouse brains. **a.** Sequence and the cloverleaf structure of mouse Arg(TCT)-4, showing the C50U mutation by an arrow. **b.** Scatter plots indicate differential abundance analysis of cyto- and mt-tRNA isodecoders from 12-week-old wild-type (B6N) and mutant (B6J) mice (n = 1). The number of cyto-tRNAs detected was 205 and 213 in WT and mutant, respectively. A total of 22 mt-tRNA isodecoders was detected in each sample. Shown in each comparison are log-transformed read counts normalized by DESeq2 and the Pearson *r*^2^ in each. **c, d.** Comparison of the normal and mutant mouse for the frequency of RT stops (pink) and of RT misincorporation (blue) at each readable modification in Arg(TCT)-4 (**c**) and in the entire tRNA pool (**d**) (n = 1).

We determined if the reduction of Arg(TCT)-4 altered its modification profile. This was to address the possibility of inter-dependence between abundance and modification of tRNA. Notably, each tRNA is modified in a defined order by modification enzymes, some acting early during the precursor state before processing, while others acting later after processing and during tRNA folding and maturation. The deficiency in processing of Arg(TCT)-4 might alter the hierarchy of its modification pathway. However, mapping and quantification of all RT-readable modifications in Arg(TCT)-4 showed a similar profile between the normal and mutant brain, while also displaying the sensitivity to detect the C50U mutation in the mutant (Fig. 5c). Likewise, mapping and quantification of all RT-readable modifications in the total tRNA pool showed a similar profile between the two (Fig. 5d). Thus, while the abundance of Arg(TCT)-4 in the mutant brain is reduced, it does not change the global tRNA modification profile, providing new insight into the genetic phenotype of the mutant mouse.

### Comparison of human and mouse tRNA modification profiles

There is currently no tRNA modification profile of the mouse genome, even though mouse is an important model organism for studies of the human genome, due to its accessibility to genetic manipulation to shed light on gene function. Our profiling of tRNA modifications of the normal mouse brain thus offered a platform to compare with the profiling of human K562 cells, providing insight into the function and evolution of the enzyme for each modification. As mutations in tRNA modification enzymes are closely linked to human diseases^60^, this comparison has the potential to uncover the underlying basis of pathology using mouse genetics.

We analyzed all isoacceptors in mouse and compared each to its human counterpart. In each isoacceptor, we interpreted a modification as the observation of RT misincorporation and mapped it to the most common chemical structure occurring at that position in the tRNA sequence framework (Fig. 2a). For example, an RT misincorporation at nucleotide G9 was interpreted as the m^1^G9 modification (Fig. 2a). We estimated the degree of each modification by the average frequency of RT misincorporation of all of its isodecoders. This average was used as the indicator of the level of the modification, based on the linear correlation between the frequency of RT misincorporation and the level of a modification (Fig. 2g). To improve confidence of the analysis, we defined modifications as RT misincorporation frequencies that are greater than 0.1 and we only analyzed the dataset for cyto-tRNAs to avoid the noise from mt-tRNAs whose non-canonical sequence frameworks complicate sequence alignment between the two organisms.

For each isoacceptor, we determined the “mouse relative to human” value using the frequency of mouse subtracted from the frequency of human. Profiling of the “mouse relative to human” value at each nucleotide position showed considerable variations across all isoacceptors, indicating the process of evolution. While most variations occurred as changes of the degree of misincorporation from mouse to human, possibly due to the sequence context, a few occurred as complete loss of a modification from one genome to the other. Notably, at position 26 that is typically modified to m^2^_2_G26 (Fig. 2a), while the modification is present in mouse Asn(GTT), it is absent from the human counterpart. Conversely, at position 27 that is typically modified to m^2^ G27, while the modification is present in Tyr(GTA) of human, it is lost in mouse. Similarly, at position 58 that is typically modified to m^1^A58, while the modification is present in Gly(GCC) of human, it is lost in mouse. In each case, the mouse and human share the same nucleotide in the isoacceptor, suggesting that the loss of the modification from one genome is a result of divergence between the two organisms. Notably, biosynthesis of m^2^_2_G26/27 is catalyzed by the same enzyme TrmT1, while that of m^1^A58 is catalyzed by TrmT61A. Neither of these enzymes is essential for life, supporting the notion that the lack of their modification products in certain tRNAs still maintains cell viability.

We then examined modifications that have demonstrated disease relevance^60^. Notably, m^1^A9 and m^1^G9 are synthesized by the enzyme TrmT10A; m^2^ G26 and m^2^ G27 are synthesized by TrmT1; m^5^C32 is synthesized by Mettl6; I34 is synthesized by the Adat2/Adat3 complex; m^1^G37 is synthesized by Trm5; and m^1^A58 is synthesized by TrmT61A. Mutations in each of these enzymes have been linked to a spectrum of diseases^60^, e.g., microcephaly, intellectual disability, neuropathy, and cancer. For each of these modifications, we performed direct comparison of the misincorporation frequency of mouse vs. human (Fig. 6b). An advantage of direct comparison is the clear identification of modifications that are absent from each organism. This information cannot be obtained by relative comparison, where a relative value of 0 could mean identical value of 0.5 in both mouse and human or a value of 0 of each. As most disease-relevant modifications are highly conserved from mouse to human, the absence of a modification suggests new biology.

**Figure 6.**
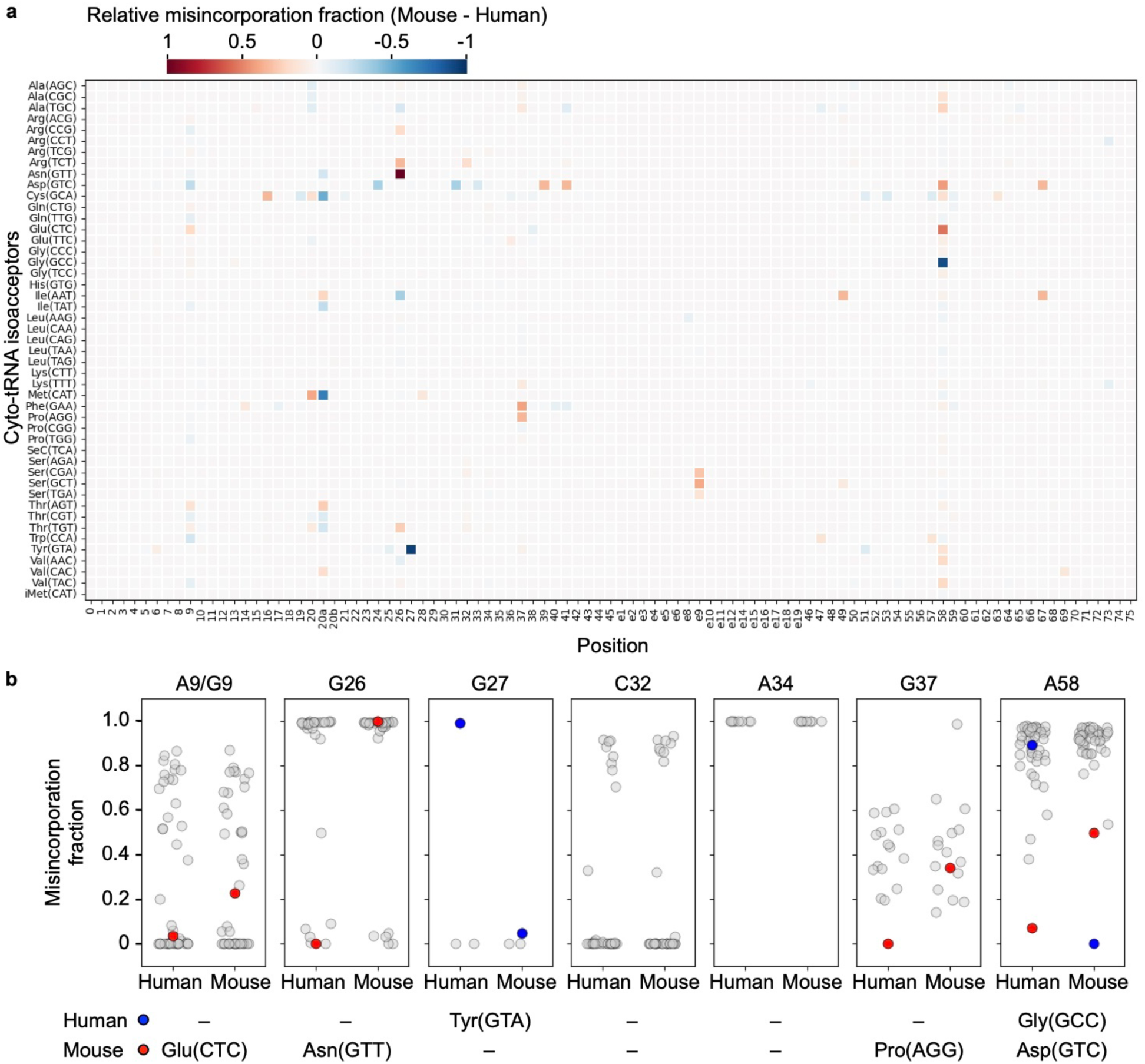
Comparison of human and mouse tRNA modification profiles. **a**. Heat map of the misincorporation frequency of WT mouse (n = 1) relative to human K562 cells (n = 4; biological replicates) for each isoacceptor (y axis) at each nucleotide position (x axis). Colors from red to blue are shown from +1 to –1 in the descending order. The misincorporation frequency of each isoacceptor is the average of the misincorporation frequencies of all isodecoders. The value of mouse relative to human is calculated by subtracting the human frequency value from the mouse frequency value. **b.** Dot plots show side-by-side direct comparison of the misincorporation frequency at A9/G9 for m^1^A9/m^1^G9 modification, at G26 for m^2^ G modification, at G27 for m^2^ G27 modification, at C32 for m^3^C32 modification, at A34 for I34 modification, at G37 for m^1^G37 modification, and at A58 for m^1^A58 modification. Blue and red dots indicate modifications that are present in human or mouse, respectively (shown in the captions below each panel).

We found that, among all tRNAs in each organism that contain A9/G9, while m^1^G9 is present in Glu(CTC) of mouse, no modification is present in the G9 nucleotide of the human counterpart (Fig. 6b). Similarly, among all tRNAs that contain G26 or G27, while m^2^ G26 is present in Asn(GTT) of mouse, it is absent from the human counterpart, although conversely, while m^2^ G27 is present in Tyr(GTA) of human, it is absent from the mouse counterpart. In contrast, among all tRNAs that contain C32, both human and mouse share the same subsets of tRNAs that possess or lack the m^3^C32 modification. Similarly, among all tRNAs that contain A34, the A-to-I conversion was uniformly observed from mouse to human, without exception. These two cases indicate a parallel development of each modification enzyme between the two organisms. Among all tRNAs that contain G37, while m^1^G37 is present in Pro(AGG) of mouse, it is absent from the human counterpart. Finally, among all tRNAs that contain A58, while m^1^A58 is present in Asp(GTC) of mouse, it is nearly absent from the human counterpart, whereas conversely, while the methylation is present in Gly(GCC) of human, it is absent from the mouse counterpart. Thus, except for m^3^C32 and I34, the implication in each case is that the modification enzyme has diverged from mouse to human.

### An unusual modification profile of human Pro(AGG) tRNA

One of the most surprising differences between mouse and human modification profiles is the presence of m^1^G37 in mouse Pro(AGG) but the absence from the human counterpart (Fig. 6b, Supplementary Fig. 4a, b). In this tRNA, where the wobble nucleotide A34 is modified to I34 in both organisms (Fig. 7a), the lack of m^1^G37 from human Pro(AGG) is unique, as the modification is present in the other two proline isoacceptors Pro(TGG) and Pro(CGG) (Fig. 7a). This absence of m^1^G37 from human Pro(AGG) is confirmed in both datasets of Induro-tRNAseq and mim-tRNAseq^34^, showing that G37 in Pro(AGG) is read as a pure G, but that it is read as a mixture of G/C/T in Pro(TGG) and Pro(CGG) (Fig. 7a). These read-identity signatures are consistent with G37 in Pro(AGG) but m^1^G37 in Pro(TGG) and Pro(CGG) (Supplementary Fig. 3c).

**Figure 7.**
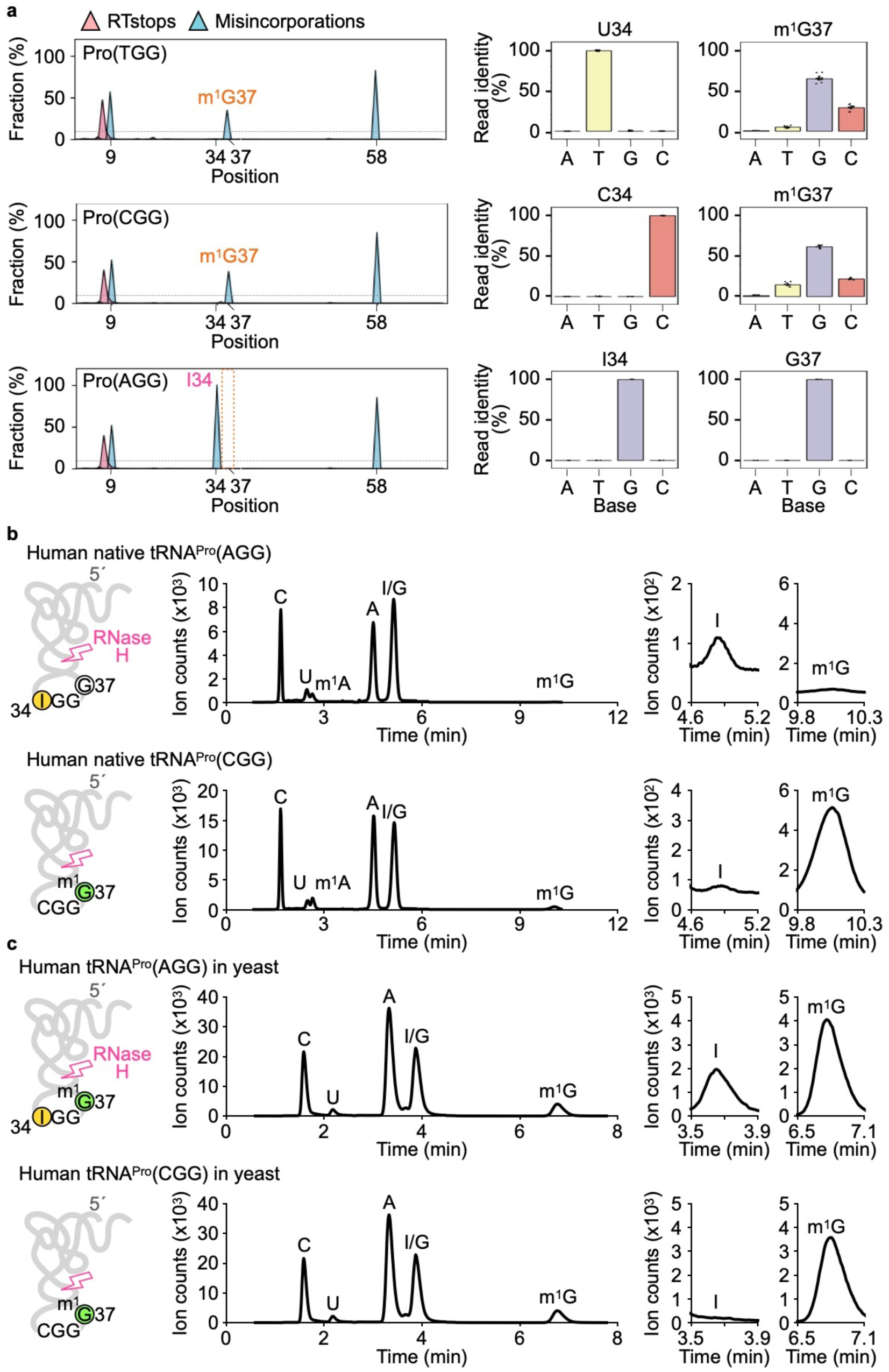
The modification profile of human proline isoacceptors. **a.** Left: The average frequency of RT stops (pink) and RT misincorporation (blue) at each readable modification in the three proline isoacceptors of K562 cells (n = 4; biological replicates). Right: Bar graphs indicate the average frequency of read identity of each isoacceptor at positions 34 and 37, respectively. Error bars indicate 95% of confidence intervals. **b.** LC-MS analysis of human Pro(AGG) and Pro(CGG) isoacceptors expressed and purified from K562 cells. Left: Isolation of the 3’-fragment of each tRNA after a targeted RNase H cleavage, where I34 is circled in yellow, G37 is circled in white, and m^1^G37 is circled in green. Middle: Ion counts (x 10^3^) of the detected nucleosides in LC-MS analysis as a function of elution time. Right: Ion counts (x 10^2^) of the detected I nucleoside at position 34 and m^1^G at position 37. **c.** LC-MS analysis of human Pro(AGG) and Pro(CGG) isoacceptors expressed and purified from yeast cells. Left: Isolation of the 3’-fragment of each tRNA after a targeted RNase H cleavage, where I34 is circled in yellow, G37 is circled in white, and m^1^G37 is circled in green. Middle: Ion counts (x 10^3^) of the detected nucleosides in LC-MS analysis as a function of elution time. Right: Ion counts (x 10^3^) of the detected I nucleoside at position 34 and m^1^G at position 37. Select chromatograms of the I and m^1^G nucleosides were amplified for better visualization of their abundance levels.

Historically, m^1^G37 has been strongly associated with proline isoacceptors. It is required to maintain the translation reading frame at proline codons^61–64^; lack of m^1^G37 induces +1 frameshifting on the ribosome, leading to cell death^61, 62, 65^. It is also required to promote efficient prolyl-charging to tRNA^Pro^; lack of m^1^G37 causes deficiency of charged tRNA^Pro^, leading to ribosome stalling and induction of a stress response similar to that of nutrient starvation^66–68^. In bacteria, which have three proline isoacceptors (TGG, CGG, and GGG), m^1^G37 is most critical for the activity of Pro(TGG), the most abundant isoacceptor that is also essential for life^68^. Loss of m^1^G37 impairs wobble reading of Pro(TGG) and fails to resolve codon usage bias^67^. In eukaryotes, the three proline isoacceptors have shifted to TGG, CGG, and AGG, where Pro(AGG) is the most abundant in species having all three isoacceptors. While the importance of m^1^G37 in eukaryotes has not been studied, its absence from the most abundant proline isoacceptor is unusual. Further, the absence of m^1^G37 from Pro(AGG) is so far only found in human, as m^1^G37 is readily detected in the isoacceptor in mouse (Supplementary Fig. 4b, 5) and in two yeast strains and in fly^34^.

We thus sought to confirm the absence of m^1^G37 from human Pro(AGG) by an LC-MS analysis, an independent method from tRNAseq. We isolated Pro(AGG) and as a control Pro(CGG) from cultured K562 cells, using a biotin-tagged oligonucleotide complementary to each in an affinity pull-down method (Supplementary Fig. 6a, Supplementary Table 3). To improve the specificity of the affinity pull-down, we used locked nucleic acid (LNA)-based oligonucleotides^69^ (Supplementary Fig. 6b, Supplementary Table 3). The homogeneity of each isolated tRNA was confirmed by Sanger sequencing (Supplementary Table 4). To enrich the nucleotides of interest in each purified tRNA and, more importantly, to eliminate the m^1^G9 present in each tRNA, we developed a method to isolate the 3’-fragment of each tRNA. We targeted RNase H to cleave each tRNA at a position 5’ to the wobble nucleotide^70^ and separated the 3’-fragment containing the anticodon loop (∼51 nucleotides) from the 5’-fragment containing m^1^G9 (∼25 nucleotides) by denaturing PAGE (Supplementary Fig. 6c). The 3’-fragment was purified from the gel and analyzed for nucleosides by LC-MS.

An LC-MS analysis of each 3’-fragment verified the lack of m^1^G37 in Pro(AGG) but its presence in the control Pro(CGG) (Fig. 7b). The identity of each fragment was confirmed by the presence of I34 in Pro(AGG) but the lack of I34 in Pro(CGG). To validate the significance of this finding, we also expressed these human isoacceptors separately in a yeast strain^71^ and isolated each by affinity-purification. Importantly, LC-MS analysis now detected the presence of m^1^G37 in the 3’-fragment of human Pro(AGG), as confirmed by its association with I34, as well as the presence of m^1^G37 in the 3’-fragment of human Pro(CGG) (Fig. 7c). Thus, the same human Pro(AGG) when expressed in K562 cells lacks m^1^G37, but when expressed in yeast contains m^1^G37. This is in complete agreement with the sequencing results in that, while m^1^G37 is absent from human Pro(AGG), it is present in the yeast counterpart. We attribute the difference between human and yeast to the divergence of the methyl transferase Trm5, responsible for m^1^G37 synthesis in eukaryotes and archaea^72–74^.

## Discussion

While RTs are critical for research and biotechnology, their mechanisms for reading RNA modifications for end-to-end cDNA synthesis are poorly understood. This problem is most serious for sequencing tRNA. Additionally, while group-II intron RTs have acquired high processivity relative to viral RTs, only two members (TGIRT and Marathon) are used in tRNA research, although their relatedness is unknown and their reading of each tRNA modification in response to temperature and time is unknown. Here we present Induro as a new group-II intron RT for genome-wide profiling of tRNA, adding it to the limited number of enzymes of the family that have been shown to successfully overcome the stable structure and extensive modifications in tRNA. We demonstrate the processivity of Induro to profile changes in the abundance and modifications of cellular tRNA in cultured and in tissue samples using Induro-tRNAseq, which is simpler, more comprehensive, and has better performances in specific cases relative to mim-tRNAseq^34^. A unique strength of this work is the mechanistic investigation of Induro for its processivity, its activity with changes of temperature and time, and its response to the sequence and structural context of each readable modification. A second is the direct comparison of Induro with TGIRT, two members of the group-II intron family, to produce two datasets for each modification to strengthen the power of prediction of the modification. A third is the new insight into how human differs from mouse in tRNA modifications, each interpreted in the context of the modification enzyme in association with human disease, thus shedding light onto tRNA evolution from mouse to human with the possibility to use mouse genetics to understand human pathology due to mutations in tRNA modification enzymes. Combined, we provide powerful new information as the engine to propel innovative tRNA research that will impact on the currently vastly expanding fields of RNA epitranscriptomics. The fundamental advance of this work is its ability to enable users to choose the best RT and the most optimal condition for the experiment at hand; to facilitate interpretation of not only the annotated but also the non-annotated modifications; to offer two misincorporation profiles for one modification to strengthen the power of prediction; and to allow new insight into genome evolution of tRNA modifications and its implications in human disease.

We show that Induro increases its processivity and readthrough of tRNA with time at 42 °C, but not at its typical temperature of 55 °C. The increased readthrough of Induro overcomes all but two modifications for end-to-end cDNA synthesis, showing strictly RT stops only at acp^3^U20 and ms^2^i^6^A37/ms^2^t^6^A37 (Supplementary Fig. 2a). Mechanistically, the increased readthrough is achieved by removal of RT stops without altering the frequency of RT misincorporation (Fig. 2d). Indeed, the frequency of RT misincorporation of Induro is constant across temperature and time (Supplementary Fig. 2b), indicating that it is an intrinsic feature that is built into the sequence and structure of the nucleotide-selection site of the enzyme. We show that the removal of RT stops is not uniform but is variable in degree with the position and chemical nature of a modification (Fig. 2d-f). While there is minor removal of RT stops at m^3^C32, I34, yW37, or ms^2^i^6^A37/ms^2^t^6^A37, a larger scale of removal of RT stops is observed at m^2^ G26/27 and at m^1^I37 (Fig. 2d). A positional effect is shown for the same modification that occurs at multiple locations, most notably for m^1^A and m^1^G. While the reduction of RT stops with time at 42 °C is minimal at position 9, which is within the tight turn of the tRNA L-shape, it is stronger at positions 37 and 58, which have fewer structural barriers (Fig. 2e, f). The sensitivity of Induro to the local structural environment of tRNA suggests that the developed RT profiles can serve as a useful reference to distinguish similar situations in other structured RNAs, such as rRNAs, lncRNAs, and regulatory RNAs, where one chemical modification can occur at multiple positions in one RNA molecule.

We provide a direct comparison of the read-identity profile of Induro and TGIRT (Fig. 4f, Supplementary Fig. 3b, c) to benchmark all RT-readable modifications. The impact is high, because while both Induro and TGIRT are both group-II intron RTs, Induro is new whereas TGIRT has been used in multiple tRNAseq methods^28, 29, 33, 34^, indicating that the information of this comparison can be extended to other TGIRT-generated datasets. Additionally, as the read identity of each RT is rooted in its intrinsic fidelity and processivity, our discovery of similarities and differences between the two is valuable. The largely similar read-identity profiles of the two RTs indicate their similar mechanism of nucleotide selection in response to most modifications, whereas their differences enable the use of two datasets to interpret one modification, which can then be verified by LC-MS analysis. This comparison provides the data for an informed decision of choosing the best RT, or both RTs, for a specific purpose. It is also useful for other applications as well. For example, many modifications in tRNA are now found in mRNA (e.g., m^1^A, m^6^A, m^7^G, m^5^C, and ψ)^75^. To predict the RT-readable m^1^A in mRNA, our comparison of the two RTs is directly applicable, whereas to predict silent modifications, we suggest that they may become readable by replacing Mg^2+^ with Mn^2+^, given that each nucleic-acid polymerase has a distinct change of misincorporation profile from one metal ion to the other.

The utility of the read-identity profile of Induro and TGIRT is exemplified in the discovery of the unusual modification profile of human proline tRNAs (Fig. 7a). The shared read identity of the two RTs provides the basis for an accurate prediction, allowing us to assign m^1^G37 to Pro(TGG) and Pro(CGG), but to assign G37 to Pro(AGG) (Fig. 7a). The unexpected lack of m^1^G37 from human Pro(AGG) is then verified by LC-MS analysis, showing that the tRNA, when expressed and isolated from human cells, lacks the methylation and contains G37, but when expressed and isolated from yeast cells, contains the methylated m^1^G37 (Fig. 7c). Thus, the read-identity profile of each modification, as we described for Induro and TGIRT, is a compelling tool that will prove valuable in new discoveries.

The absence of m^1^G37 from human Pro(AGG) suggests intriguing biology. This absence is unique to human Pro(AGG), even at the resolution of isodecoders. Indeed, we have analyzed all isodecoders that contain both A34 and G37 across 5 species that are well separated in evolution (human *H. sapiens*, mouse *M. Musculus*, fly *D. melanogaster*, yeast *S. cerevisiae*, and yeast *S. pombe*) based on data generated in this work and in mim-tRNAseq^34^ (Supplementary Fig. 7). We find that all these isodecoders are members of three families: Pro(AGG), Leu(AAG), and Arg(ACG), and that, while A34 is uniformly modified among them, indicative of I34 at 100%, the absence of m^1^G37 is only found in Pro(AGG) of human, whereas the modification is present in all other species. Thus, the consistent lack of m^1^G37 from human Pro(AGG) is remarkable, indicating that it is a recent development that is restricted to only the most complex mammals. Even among human tRNAs, the specific lack of m^1^G37 from Pro(AGG), but not from Leu(AAG) or Arg(ACG), is interesting, suggesting a unique evolution of m^1^G37 with the proline family. Notably, in the simplified tRNA pool of *E. coli*, where each amino acid has only isoacceptors but virtually no isodecoders, while m^1^G37 is present in all isoacceptors of Leu and Pro and in the Arg(CCG) isoacceptor and is required for efficient charging of Pro isoacceptors and the Arg(CCG) isoacceptor ^66^, it is specifically associated with the proline family^67^. This association is identified in a genome-wide genetic screen for suppressors that overcome the deficiency in tRNA charging due to lack of m^1^G37. While several suppressors were found in this screen, all were mapped to the *proS* gene required for prolyl-charging to tRNAs, but none were mapped to the *leuS* or *argS* gene^67^, emphasizing the strong association of m^1^G37 with proline tRNAs in evolution.

Given the long-standing history of m^1^G37 with proline tRNAs, the specific absence of the modification from human Pro(AGG), but not from human Pro(CGG) or Pro(TGG), is highly unusual. It raises two important points for consideration. First, it suggests that the methyl transferase Trm5 for synthesis of m^1^G37 in human^73^, has evolved with unique sequence motifs that specifically discriminate against I34, whereas the counterpart yeast and mouse enzymes lack such motifs. Thus, while human Pro(AGG) when expressed in human is discriminated by human Trm5 from m^1^G37 methylation, it is accepted when expressed in yeast by yeast Trm5 for synthesis of m^1^G37 (Fig. 7c). Further biochemical and structural analysis of the human Trm5 enzyme, together with studies of yeast or mouse Trm5, will be necessary to elucidate the human-specific motifs. Second, the lack of m^1^G37 from human Pro(AGG) suggests that the G37-bearing tRNA must function properly in protein synthesis. As shown in our work with m^1^G37 in bacterial proline isoacceptors, the methylation is required for efficient prolyl-charging of tRNA^66, 67^ and for accurate maintenance of the translational reading frame at proline codons^61–64^. How each of these important functions is achieved despite the lack of m^1^G37 from human Pro(AGG), and how the deficiency might be compensated by the presence of I34, are key questions to address. Regardless of the outcome of these questions, we show here that the development of Induro-tRNAseq, coupled with mechanistic analysis of the RT, is fundamental to a new frontier of tRNA research in human biology.

## Methods

Unless otherwise noted, kit-based protocols described below followed the manufacturer’s instructions of each.

### Cell culture

Human embryonic kidney HEK293T cells (ATCC, #CRL-3216) and chronic myeloid leukemia K562 cells (ATCC, #CL-243) were grown at 37 °C with 5% CO_2_ in Dulbecco’s Modified Eagle’s Medium (DMEM) (Corning) with 10% fetal bovine serum (Corning). *S. cerevisiae* BY4741 strain (ATCC, #201388) was grown at 30 °C in yeast synthetic dropout medium lacking leucine^71^.

### RNA extraction and tRNA isolation

Total RNA from HEK293T and K562 cells was extracted with TRIzol (Invitrogen) according to the manufacturer’s instructions. Total RNA from the cerebellum of 12-week-old C57BL/6J (B6J) and congenic B6J.B6N*^nTr^*^20^ mice was isolated as previously described^59^. The mouse brains used in this study were obtained from animals that were maintained under the institutional IACUC guidelines, compliant with those of the American Veterinary Medical Association for the husbandry and euthanasia. The tRNA pool was isolated from total RNA by 12% denaturing PAGE/7 M urea in TBE (90 mM Tris-borate pH 8.3 and 2 mM EDTA) as gel slices, which were crushed with a disposable pestle in tRNA extraction buffer (0.3 M Na Acetate (NaOAc), pH 5.0, 0.25% SDS, and 1 mM EDTA) as described^34, 54^. The tRNA pool was filtered through a Costar Spin-X 8163 centrifuge tube filter (Corning), precipitated with isopropanol in the presence of 20 µg glycogen (RNA grade; Thermo Scientific), and dissolved in RNase-free water. Approximately 12% of the input total RNA was recovered as tRNA. To compare with mim-tRNAseq dataset^34^, Induro-tRNAseq was performed with 4 biological replicates of total RNA from K562 cells, 2 biological replicates of gel-purified tRNA pool of K562 and HEK293T cells, and 2 biological replicates of total RNA of HEK293T cells. Only one biological sample was obtained from WT or mutant mouse brain tissues.

### tRNAseq library preparation

Prior to starting the workflow, the integrity of each sample (500 ng of total RNA or 125 ng of gel-purified tRNA pool) was verified by 12% denaturing PAGE/7 M urea. For the Induro time course experiment, the workflow was carried out twice to provide two technical replicates using one of the four biological replicates of total RNA of K562 cells. The tRNA in each sample was first deacylated by incubation for 45 min at 37 °C in 10 µL 75 mM glycine, pH 9.0, with 1 U/µL Superase-In (Thermo Fisher). The reaction was diluted to 100 µL with 1X T4 PNK (polynucleotide kinase) buffer and incubated for 30 min with 1 µL T4 PNK (10 U/µL; NEB M0201) to remove 3’ phosphoryl groups. The deacylated RNA was ethanol precipitated with 20 µg of glycogen and the pellet was dissolved in 7.0 µL water.

T4 Rnl2 (NEB M0239L)) was used to catalyze splint-mediated 3’ ligation of each tRNA library to a barcoded 35-mer RNA/DNA adaptor bearing a 5’-phosphate and a 3’-amino group (oligonucleotides listed in Supplementary Table 1). Typically, 4 ligation reactions were carried out in parallel with each in 20 µL and containing a separate tRNA library, a uniquely barcoded 3’-adaptor (0.25 µM), a corresponding DNA splint (0.32 µM), 0.5 U/µL T4 Rnl2, and 1 U/µL Superase-In RNase inhibitor (Thermo Fisher Scientific) in a buffer containing 400 µM ATP, 10 mM MgCl_2_, 1 mM DTT, 10% PEG8000, and 50 mM Tris-HCl, pH 7.5. Reactions were briefly heat-cooled at 60 °C for 2.5 min prior to adding T4 Rnl2 and kept at 16 °C for 45 min and then 25 °C for 45 min. Ligation was terminated by combining all 4 reactions into 20 µL 0.1 M EDTA, followed by extraction with phenol-chloroform-isoamyl alcohol (25:24:1), pH 5.2, and purified through a Zymo Oligo Clean & Concentrator cartridge (Zymo Research). The nucleic acid in the multiplexed library was eluted in 11 µL of RNase-free water.

Each library was converted to cDNA with Induro RT (NEB M0681S) using a 57-mer ssDNA as a primer. The sequence of the primer was 5’-<u>pRN</u>AGATCGGAAGAGCGTCGTGTAGGGAAA-GAG/iSp18/GTGACTGGAGTTCAGACGTGTGCTC-3’, where p is a phosphate, R is purine, N is any nucleotide, and iSP is an 18-carbon chain spacer. Each RT reaction was performed in 20 μL with 0.5 µM RT primer, 0.75 mM dNTPs, 10 mM DTT, 1 U/µL Superase-In, and 10 U/µL Induro RT in 1X Induro buffer. The reaction was briefly heat-cooled at 60 °C prior to adding the RT and was incubated overnight (12-16 h) at 42 °C. After cDNA synthesis, the RNA was hydrolyzed by adding 1 µL of 5 M KOH and heating at 90-95 °C for 3 min. The solution was neutralized with 0.25 M NaOAc, pH 5.0, extracted with phenol-chloroform-isoamyl alcohol (25:24:1), and the cDNA precipitated with 3 volumes of ethanol. The pellet was dissolved in 8 µL 7 M urea-TBE with 0.01% xylene cyanol and bromophenol blue. After a 2 min heat-cool at 80-85 °C, the cDNA was loaded onto a denaturing 10% PAGE/7 M urea gel (8 x 7 x 0.1 cm) and electrophoresed at 200 V in hot TBE for 20 min. Included in the gel as controls were an RT reaction conducted in the absence of tRNA and a Small Range RNA Ladder (NEB N0364S). The gel was stained with SYBR gold and visualized using a ChemiDoc imager (BioRad). The gel was excised in the range of 90-180 nucleotides (nts) to include tRNA templated full-length and truncated cDNA. cDNA was extracted into 500 µL TE by incubation in a MultiTherm Shaker (Benchmark Scientific) at 70 °C/1500 rpm for 1-2 h. After gel removal, the cDNA was precipitated with isopropanol in the presence of 20 µg glycogen carrier and the pellet dissolved in 5.5 µL water.

The cDNA was circularized by incubation with Circligase (Lucigen) for 3 h at 60 °C in a 10 µL reaction as described^54^. The reaction was terminated by heat inactivation (at 80 °C for 10 min) and stored at –20 °C until PCR amplification.

The circularized cDNA (1 µL) was amplified by PCR using Q5 Hot Start High-Fidelity 2X Master Mix (NEB M0494L) in a 25 µL reaction containing 12.5 pmoles of forward and reverse primers (primers listed in Supplementary Table 2). A unique barcoded reverse primer was used for each group of 4 multiplexed cDNA pools. The thermocycler was programmed for 30 sec at 98 °C, followed by at least 5 cycles of 10 sec at 98 °C, 10 sec at 62 °C, and 10 sec at 72 °C. To determine the optimal cycle number, separate reactions were terminated after 5, 6, 7, and 8 cycles. After clean-up by ethanol precipitation or passage through a Zymo Oligo Clean & Concentrator cartridge, these reactions were electrophoresed on a non-denaturing 8% PAGE gel alongside O’RangeRuler 10 bp DNA and GeneRuler 50 bp DNA ladders (Thermo Fisher), followed by SYBR gold staining. The dsDNA product (150-240 bp in length) with the highest yield but without contaminating higher MW DNA was excised from the gel and eluted into 500 µL TE by continuous mixing at room temperature overnight. After clarification of the suspension through a Costar Spin-X cartridge, the DNA was ethanol precipitated with 20 µg glycogen and dissolved in 20 µL TE. The average yield of gel-purified DNA from each multiplexed sample of 4 was approximately 50 ng as determined by Qubit (Invitrogen) and Bioanalyzer (Agilent) analysis.

### Sequencing

Equimolar amounts of the PCR-amplified cDNA libraries were pooled and loaded onto the Illumina NextSeq 500 platform and a 2×75 paired end sequencing run was performed.

### Synthesis of mt-Leu(TAA) for calibration analysis

The 5’-RNA oligonucleotide of mt-Leu(TAA) was chemically synthesized encoding the sequence from nucleotides 1 to 17, containing either G9 or m^1^G9. The 3’-RNA oligonucleotide of the tRNA was transcribed by T7 transcriptase and gel purified. The 5’- and 3’-RNA oligonucleotides were mixed together in equal molar concentration, heat-cooled, and covalently joined by T4 Rnl2. The reconstituted full-length mt-Leu(TAA) was separated from the component RNAs by a denaturing PAGE/7 M urea gel and extracted from the gel. The concentration of the reconstituted full-length tRNA was determined. For calibration analysis, the G9-containing and the m^1^G9-containing full-length tRNAs were mixed in fractions of 1:0, 3:1, 1:3, and 0:1, where the methylated tRNA represented 0, 25, 75, and 100%.

### Expression of human Pro(AGG) and Pro(CGG) in yeast

Each human tRNA gene was inserted into a *S. cerevisiae* tandem tRNA operon, downstream from the Arg(TCT) gene, replacing the Arg(GTC) gene as described^76^. The tandem tRNA operon sequence was chemically synthesized as a gBlock DNA fragment by IDT (Integrated DNA Technologies), with a BamHI restriction site on the 5’-end and a HindIII restriction site on the 3’- end. The gBlock DNA and a pRS425 high copy number plasmid^77^ were digested with HindIII (NEB R0104S) and BamHI (NEB R0136S) and ligated together with T4 DNA Ligase (NEB M0202S). The plasmid was then introduced into bacteria by transformation, purified, and subsequently introduced into the *S. cerevisiae* BY4741 strain by transformation. Expression of each human tRNA gene in yeast was verified using a label-free assay for aminoacylation of tRNA as described^78^.

### LC-MS analysis of m^1^G37 in human Pro(AGG) and Pro(CGG)

Total RNA from K562 cells was extracted with TRIzol (Invitrogen) according to the manufacturer’s instructions. Total RNA was extracted from yeast by resuspension of cells in 15 mL of Buffer A (50 mM NaOAc, pH 5.0, and 10 mM Mg(OAc)_2_), followed by shaking at 37 °C for 1 h with 2 volumes of phenol. Shaking was continued for 30 min after adding 0.1 volume of chloroform. Phases were separated by centrifugation at 6500 rpm for 20 min at 4 °C. The aqueous phase was collected, and RNA precipitated in 2.5 volumes of ethanol and 0.1 volume of 2.5 M NaOAc pH 5. The tRNA pool was isolated from yeast and human total RNA as described^79^. Native tRNAs of interest were affinity purified from total tRNA using biotinylated oligonucleotide probes (IDT) immobilized to a streptavidin sepharose (Cytiva) solid support^80^. To enhance the specificity of affinity purification, the biotinylated oligonucleotide was synthesized with a chimeric DNA-LNA backbone targeted to hybridize to I34 and G37 in Pro(AGG), and targeted to hybridize to G30 and C34 in Pro(CGG). Sequences of biotinylated oligonucleotides are listed in Supplementary Table 3. Hybridization of tRNA to the biotinylated oligonucleotide was performed in 10 mM Tris-HCl, pH 7.5, 0.9 M TEA-Cl, 0.1 mM EDTA, at 42 °C. After extensive washes, the tRNA was eluted from the solid support by two consecutive 5 min incubations at 65 °C in TE (10 mM Tris-HCl, pH 8.0, 1 mM EDTA).

To verify purity, each affinity-purified tRNA was processed for Sanger sequencing. Briefly, 2.4 pmole of each affinity-purified tRNA was mixed with 5 pmole of a DNA primer (IDT), which was complementary to the 3’-end of the tRNA and had a 3’-overhang (Supplementary Table 4), and the mixture was incubated at 65 °C for 5 min. This was followed by addition of 1 μL M-MLV RT (Promega) to initiate reverse transcription in a 20 μL reaction of the M-MLV buffer, with Ribolock RNase Inhibitor (Thermo Scientific) and 1 mM dNTP. After incubation at 25 °C for 5 min, 42 °C for 1 h, and finally 70 °C for 5 min, the synthesized cDNA was mixed with the forward and reverse primers (IDT) in 2X Optima Hotstart ReadyMix (FastGene) and PCR amplified. Sanger sequencing (Genewiz) confirmed the homogeneity of each affinity-purified tRNA (data not shown). To examine the methylation status only at position 37, the purified tRNA was treated with In-house purified RNase H in the presence of a chimeric DNA-2’-OMe-RNA oligonucleotide designed according to previous studies^70, 81, 82^ to direct the cleavage at or near position 26 in the tRNA (Supplementary Table 3). This resulted in two fragments, the 5’-fragment of ∼25 nts containing m^1^G9, and the 3’-fragment of ∼51 nts containing G37 or m^1^G37 (Supplementary Fig. 6a). To perform the RNase H cleavage reaction, 2 pmoles of purified tRNA were first mixed with 8 pmoles of the chimeric oligo in 3.5 µL of water, incubated at 85 °C for 10 min, and left on a bench at room temperature for 10 min. The sequence of the oligonucleotide is listed in Supplementary Table 3. After adding 10X RNase H reaction buffer and 2 pmoles of an in-house purified RNase H, the resulting 5 µL solution was incubated for 20 min at 37 °C, quenched by adding 1 µL of 0.5 M EDTA pH 8.0, and electrophoresed through a 12% denaturing PAGE/7 M Urea gel in TBE. The 3’-fragment was eluted into tRNA extraction buffer, precipitated, and dissolved in water as described earlier. Prior to nuclease digestion to single nucleosides, the 3’-fragment was incubated for 5 min at 95 °C and then cooled on ice for 3 min, followed by digestion to ribonucleotides by incubation for 2 h at 45 °C with 0.15 U of Nuclease P1 (NEB M0660S) in 10 mM ammonium acetate pH 5. Dephosphorylation was performed by adding 10X Antarctic phosphatase buffer and 0.5 µL of Antarctic Phosphatase (NEB M0289S) prior to incubation for 2 h at 37 °C. The digested mixture of nucleosides was purified away from proteins by centrifugation through a 10,000-Da MWCO spin filter (Amicon).

LC-MS analysis of nucleosides dissolved in LC/MS grade water was carried out as described^83^ on an Agilent 1260 Infinity II HPLC coupled to an Agilent 6470 triple quadrupole mass spectrometer in positive ion mode, using the dynamic multiple reaction monitoring method. Samples were run through a Hypersil GOLD aQ C18 Polar Endcapped HPLC column (Thermo Fisher, 25303-152130; 3-µm particle size, 175-Å pore size, 2.1 × 150 mm) at a flow rate of 0.4 mL/min for 14 min in 0.1% formic acid in water at a temperature of 37 °C. The parameters used for the mass spectrometer were as follows: gas flow at 12 L/min, nebulizer pressure at 20 psi, sheath gas temperature at 325 °C, and capillary voltage at 2,500V. Parent ion MS1 to deglycosylated base ion MS2 transition for each nucleoside was set as follows: *m*/*z* 268 → 136 for A, *m*/*z* 244 → 112 for C, *m*/*z* 284 → 152 for G, *m*/*z* 269 → 137 for I, *m*/*z* 282 → 150 for m^1^A, *m*/*z* 298 →166 for m^1^G, and *m*/*z* 245 → 113 for U. A standard curve of nucleosides ranging in concentrations from 5 ng/mL to 500 ng/mL for canonical nucleosides and 0.5 ng/mL to 50 ng/mL for modified nucleosides was used for quantitative analysis of I and m^1^G levels. Quantification of the nucleosides was performed using Agilent Quant-My-Way software.

### Data analysis

Read processing was performed as described^34^. Paired-end reads were merged using PEAR v0.9.6 (ref ^84^), which were then demultiplexed using cutadapt v2.5 as described^34^ and a fasta file of the first 10 nts for the different 3’-barcoded adaptors was created. Indels in the alignment to the adaptor were removed with --no-indels. The two 5’-RN nucleotides introduced by reverse transcription (see *Library Preparation* above) were trimmed from reads with *-u* 2. Reads shorter than 10 nts were discarded by cutadapt according to the parameters in -*m* 10. Processed reads of human samples were mapped to tRNA reference transcripts derived from human genome hg38 and those of mouse samples mapped to the reference derived from the mouse genome mm39 using mimseq v1.2. Mapping was as described^34^ (github.com/nedialkova-lab/mim-tRNAseq), where matured and processed tRNA sequences were mapped to MODOMICS entries using BLAST and clustered with the --*cluster* parameter using a user-defined sequence identity threshold. After clustering, reads were aligned using GSNAP to the representative cluster sequences of mature tRNAs. The following mapping parameters were used:

*H. sapiens*: --species Hsap --cluster --cluster-id 0.95 --min-cov 2000 --max-mismatches 0.1 --remap --remap-mismatches 0.075
*M. musculus*: --species Mmus --remap --remap-mismatches 0.075

Mapping rates were calculated by the number of uniquely mapped reads relative to the sum of the number of unmapped reads, multiple-mapped reads, and uniquely mapped reads. Additional quantification and statistical analysis were also based on methods described^34^. Fractions of incomplete 3’-ends (3’-N, 3’-NC and 3’-NCC, where N = the discriminator nucleotide at position 73) relative to all 3’-ends (including 3’-NCCA) were calculated per unique tRNA sequence. For differential tRNA abundance analysis, read counts of cyto-tRNAs and mt-tRNAs were normalized with DESeq2 (ref ^85^) separately and the Pearson correlation coefficient (*r*) was calculated. Rates of misincorporation were calculated by summing up counts of mismatches for all four nucleotides relative to total read counts at the position of interest. Frequency of stops was determined by dividing the number of stops at a given position by the total number of reads at that position. Readthrough at a position was obtained by subtracting the ratio of RT stops at that position from 1.0. For a given modification, RT stops were reported as the maximum value observed in a 3-nt window (–1, 0, and +1) centering on the modification to reduce the likelihood of over-estimation. Modifications in human tRNAs at known positions (9, 20, 26, 32, 34, 37, and 58), as well as non-annotated positions, were identified by RT stop or RT misincorporation of >10%. Annotated modifications were as defined^34, 56^. All data analysis was performed with in-house python written codes using Python v3.7.0. Analysis of the mim-tRNAseq dataset was performed using the following parameters:

*H. sapiens*: --species Hsap --cluster --cluster-id 0.95 --min-cov 2000 --max-mismatches
0.1 --remap --remap-mismatches 0.075
*S. cerevisiae*: --species Scer --cluster --cluster-id 0.90 --min-cov 2000 --max-mismatches
0.1 --remap --remap-mismatches 0.075
*S. pombe*: --species Spom --cluster --cluster-id 0.95 --min-cov 2000 --max-mismatches
0.1 --remap --remap-mismatches 0.075
*D. melanogaster*: --species Dmel --cluster --cluster-id 0.95 --min-cov 2000 --max-mismatches 0.1 --remap --remap-mismatches 0.075

### Data sources

The raw FASTQ files of mim-tRNAseq^34^ of wild-type human cells (HEK293T and K562), yeast (*Saccharomyces cerevisiae and Schizosaccharomyces pombe*), and fly (*Drosophila melanogaster*) were retrieved from Gene Expression Omnibus (GEO): GSE152621.

### Data availability

All sequencing data generated in the tRNA-seq experiments are publicly available through the sequence read archive (SRA) database of NCBI: PRJNA996215.

### Code availability

Scripts for data processing and figure creation are available at https://github.com/YaMing-Hou-lab/Induro-tRNAseq.

## Acknowledgements

We thank Danny Nedialkova and Andrew Behrens of Max Planck Institute at Martinsried, Germany, for discussions of mim-tRNAseq and the associated computational tool kit. We thank Tao Pan of U. Chicago and Satoshi Kimura of Harvard Medical School for support of our earlier tRNAseq. We thank Susan Ackerman and Mridu Kapur of U. California, San Diego, for providing mouse brain tissues and Tom Christian of the Hou lab and Nathan Yu and Xuemeng Sun of the Kleiner lab for assistance in LC-MS analysis of inosine and m^1^G in tRNA. This work was supported by NIH R35 GM149336 to J.P. and NIH R35 GM134931 to Y.M.H.

## Author contributions

Y.N., H.G., S.M., E.Y. and Y.M.H. conceived this study. H.G., E.Y., N.N. and Y.M.H. coordinated the collaboration. Y.N., H.G. and K.K. acquired data. N.S.L. and J.A.P. generated synthetic m^1^G9-containing RNA oligonucleotides. Y.N., H.G. and Z.S. analyzed and interpreted data. Z.S. and Y.N. performed the first pass analysis and data indexing. Y.N. performed RT profiling analysis, tRNA abundance analysis, read-identity analysis, RNase H analysis, and designed and implemented scripts for data analysis. H.M. performed affinity purification of tRNA and Sanger sequencing analysis. H.M. and R.K. acquired data for LC-MS analysis. Y.N., H.G. and H.M. prepared figures, while Y.N., H.G. and Y.M.H. prepared the manuscript.

## Additional information

Supplementary information is available online.

