## Supplemental Information for "Genome-Wide Profiling of tRNA Using an Unexplored Reverse Transcriptase with High Processivity"

**Integrated Supplementary Information**

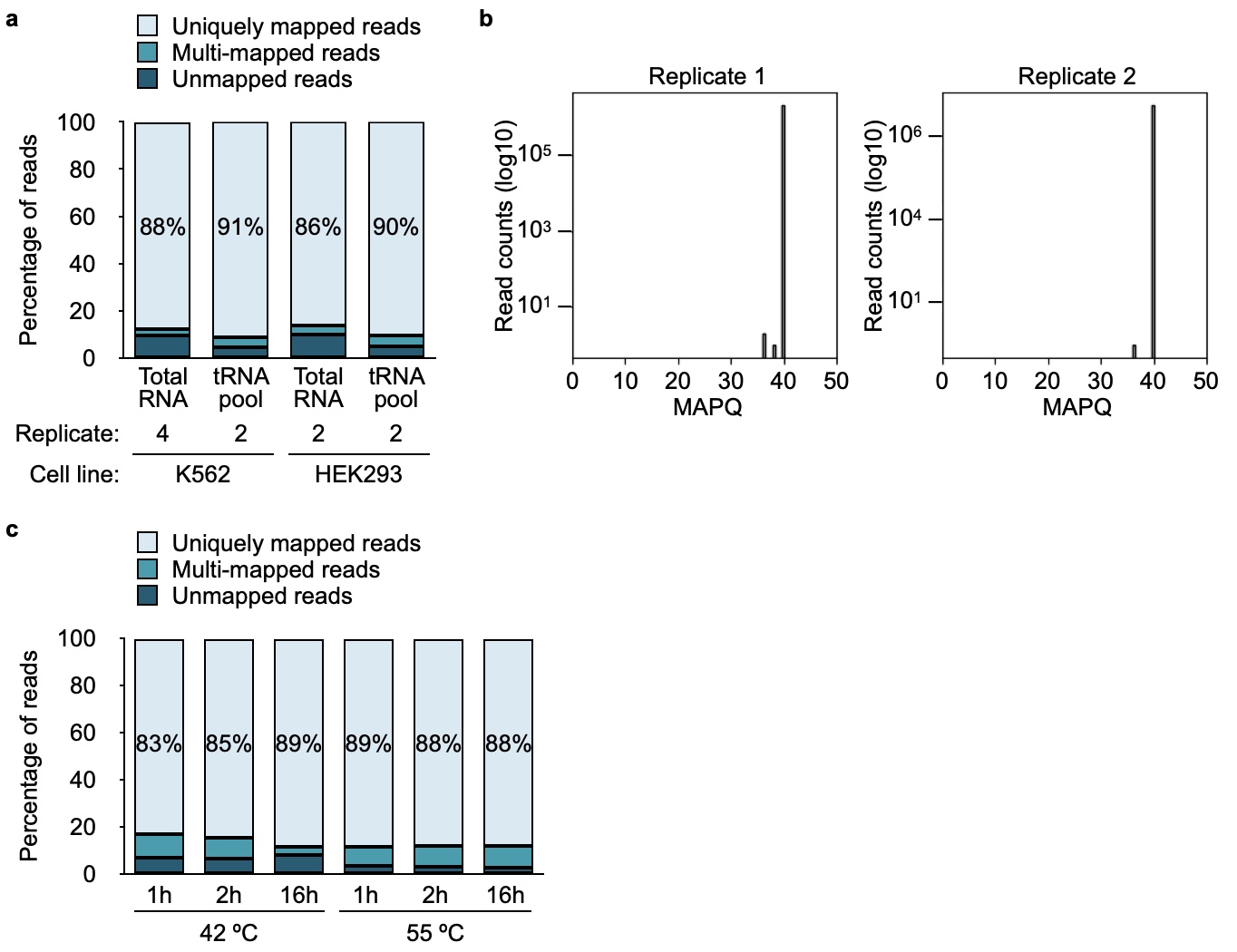

**Supplementary Figure 1. Quality analysis of Induro-tRNAseq**.

**a.** Cumulative bar graphs indicate the average mapping frequency (%) of total reads from sample preparation starting with total RNA or a purified tRNA pool of K562 and HEK293T cells. **b.** The distribution of the mapping quality (MAPQ) score in two replicates of sample preparation starting with total RNA from K562 cells. The Y-axis indicates log-transformed read counts. The coefficient of determination (*r^2^*) in each replicate is 0.9999. **c.** Cumulative bar graphs indicate the average mapping frequency (%) of total reads from sample preparation starting with total RNA of K562 cells collected at 42 ºC or 55 ºC after the RT reaction for 1, 2, or 16 h (n = 2; technical replicates).

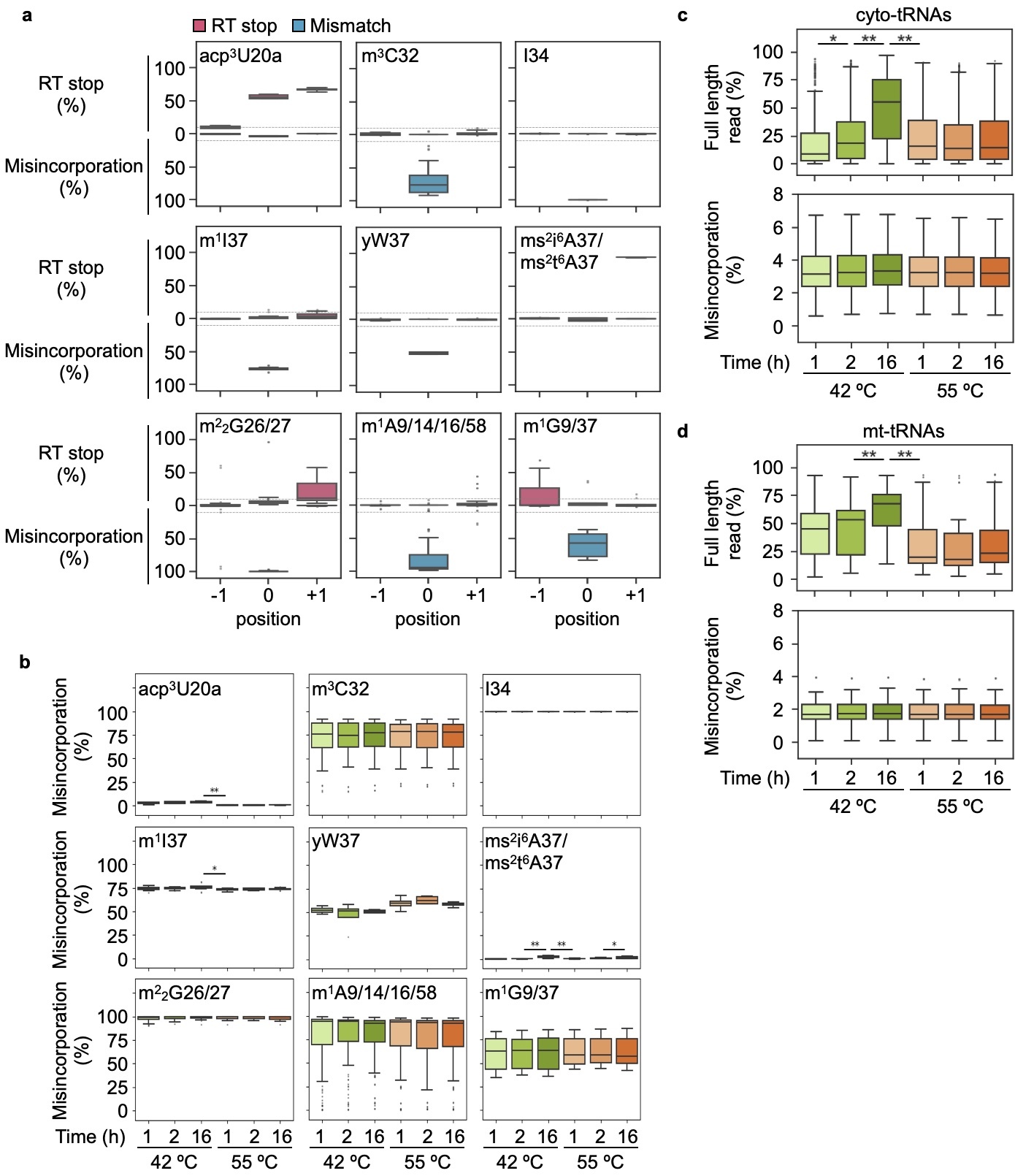

**Supplementary Figure 2.** **Profiling of readthrough of Induro reactions.**

**a.** The boxplots indicate the frequency (%) of RT stops (pink) and of RT misincorporation (blue) in a 3-nucleotide window centering on position 0 as the annotated modification. RT reaction was performed at 42 ºC for 16 h. **b.** The boxplots indicate the average frequency (%) of misincorporation at each readable modification analyzed for the RT reaction at 42 ^o^C or 55 ^o^C over the indicated time. **c.** The boxplots indicate the frequency of full-length read and of the RT misincorporation in cyto-tRNAs analyzed for the RT reaction at 42 ^o^C or 55 ^o^C over the indicated time. **d.** The boxplots indicate the frequency of full-length read and of the RT misincorporation in mt-tRNAs analyzed for the RT reaction at 42 ^o^C or 55 ^o^C over the indicated time. All samples were collected from total RNA of K562 cells as the starting material (n = 2; technical replicates). Center line: median; box limits: upper and lower quartiles; whiskers: 1.5X interquartile range; points: outliers. Student's *t*-test was performed by a two-sided analysis (**p* < 0.1, ***p* < 0.01).

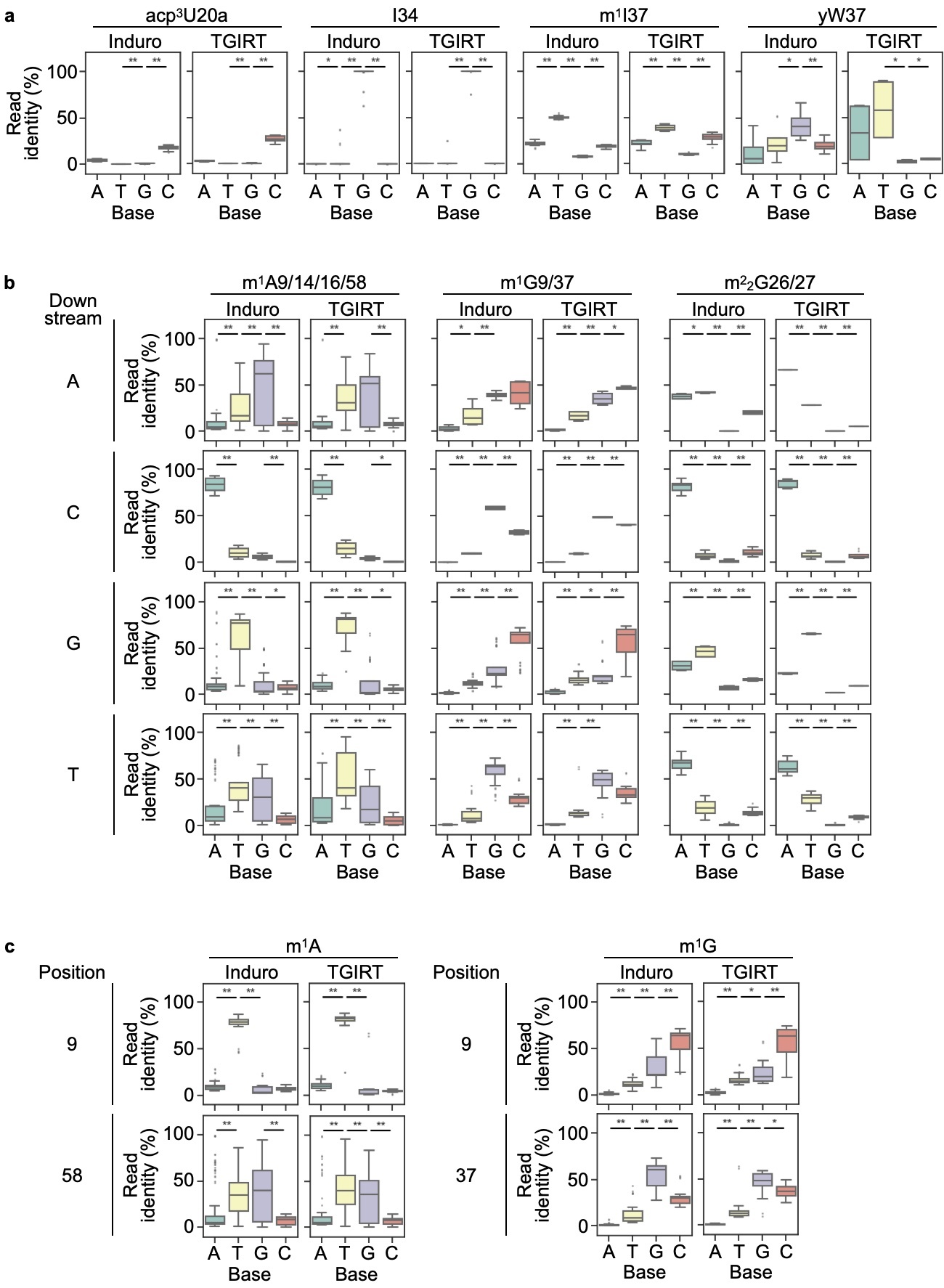

**Supplementary Figure 3. The read identities of RT-readable modifications.**

**a.** Boxplots of the read identity (%) of each context-independent RT-readable modification generated by Induro in Induro-tRNAseq (n = 4, biological replicates) and by TGIRT in datasets generated in mim-tRNAseq^34^. **b.** Boxplots of the read identity (%) of each context-dependent RT-readable modification generated by Induro and by TGIRT in response to the downstream nucleotide. **c.** Boxplots of the read identity (%) of m^1^A9 vs. m^1^A58, and of m^1^G9 vs. m^1^G37 generated by Induro and by TGIRT. All data were analyzed for samples collected from K562 cells. Center line: median; box limits: upper and lower quartiles; whiskers: 1.5X interquartile range; points: outliers. Student's *t*-test was performed by a two-sided analysis (**p* < 0.1, ***p* < 0.01).

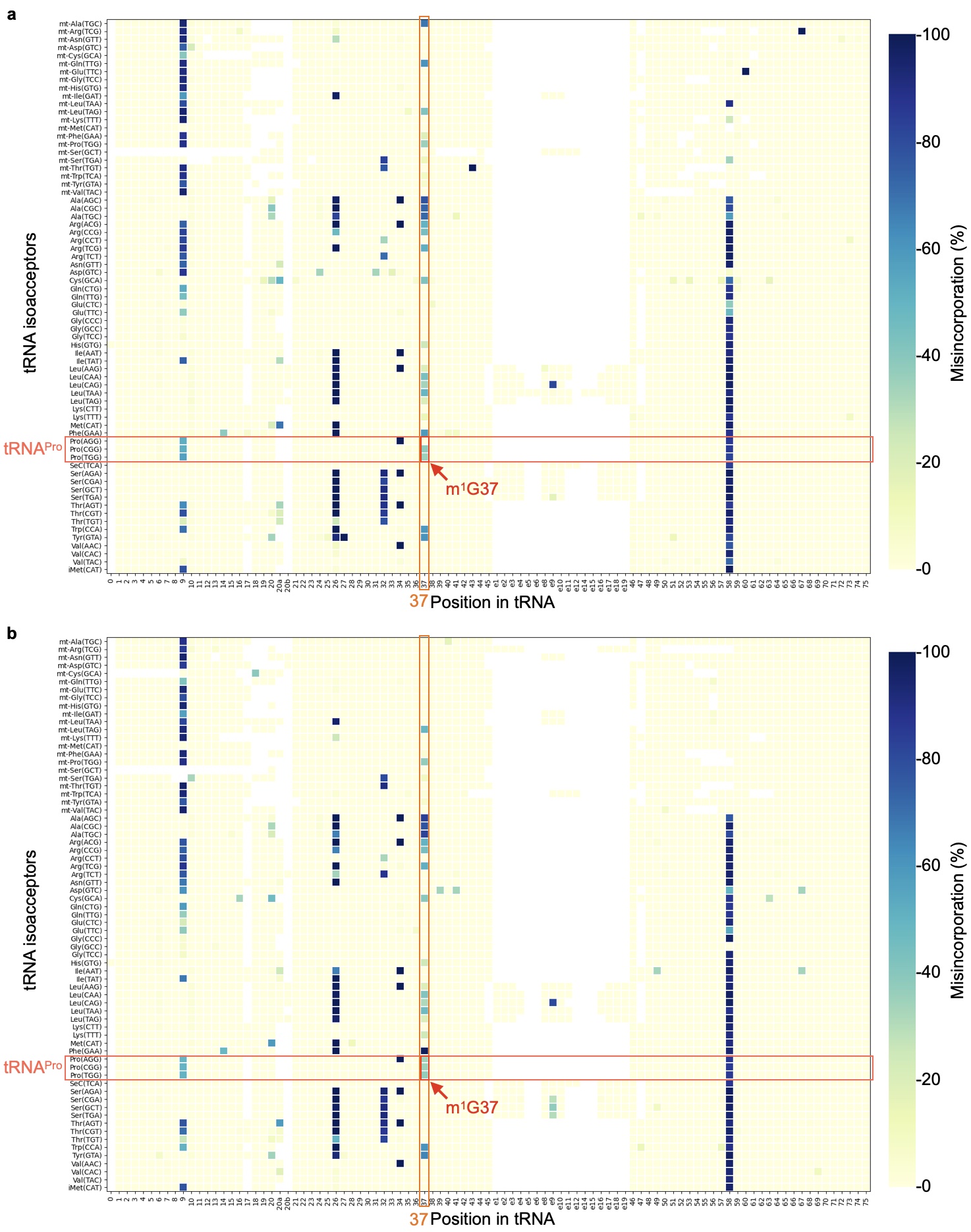

**Supplementary Figure 4.** **Profiling of tRNA modifications in human and mouse cells**.

**a.** Heatmap of the frequency (%) of RT misincorporation at each readable modification averaged from all tRNA isoacceptors collected from K562 cells (n = 4; biological replicates). Position 37 and the three proline isoacceptors are highlighted by red boxes. Note that Pro(AGG), which contains I34, does not elicit misincorporation at position 37, suggesting that the guanosine at this position is not *N*^1^-methylated. **b.** Heatmap of the frequency (%) of RT misincorporation at each readable modification averaged from all tRNA isoacceptors collected from wild-type B6N mouse brain (n = 1). Position 37 and the three proline isoacceptors are highlighted by red boxes.

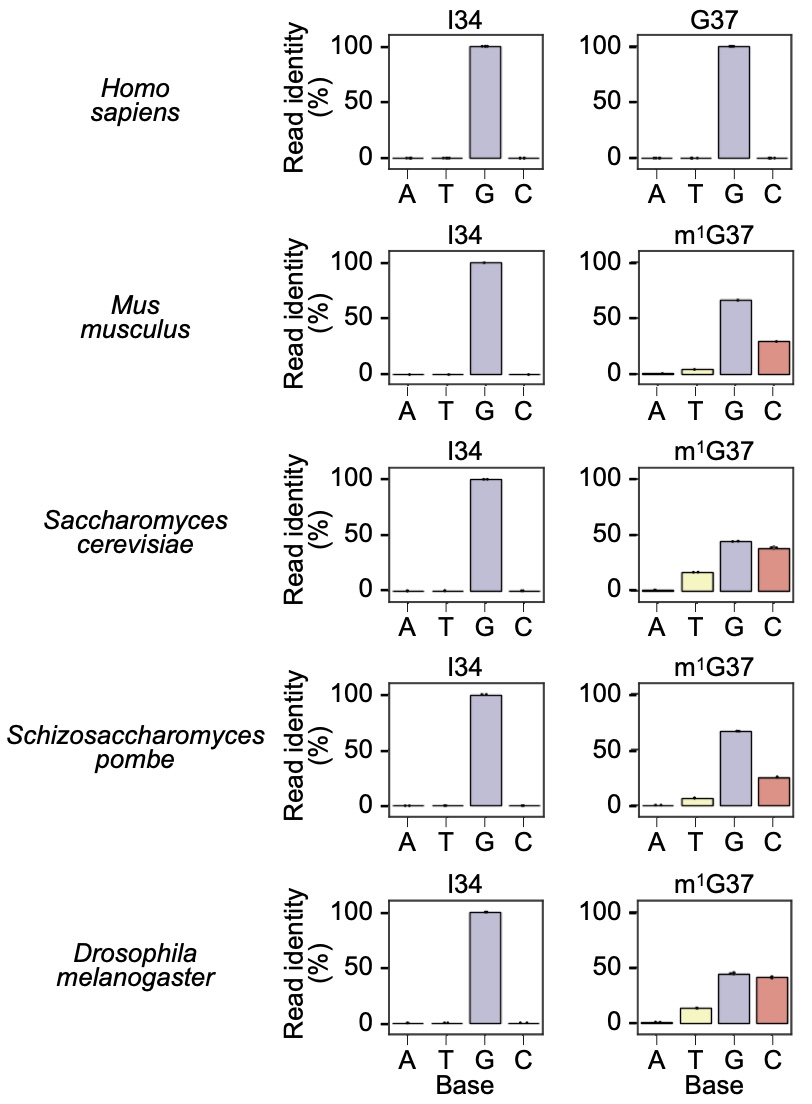

**Supplementary Figure 5.** **Specific** **absence of m^1^G37 from human Pro(AGG).**

Bar graphs showing the read identity (%) of A34 and G37 are shown for Pro(AGG) in *Homo sapiens*, *Mus musculus*, *Saccharomyces cerevisiae*, *Schizosaccharomyces pombe*, and *Drosophila melanogaster* . The data *M. musculus* is from Induro-tRNAseq of this work (n = 1), while others shown here are extracted from the published datasets of mim-tRNAseq^34^. The error bars indicate 95% of confidence intervals. Individual data points are shown in dots.

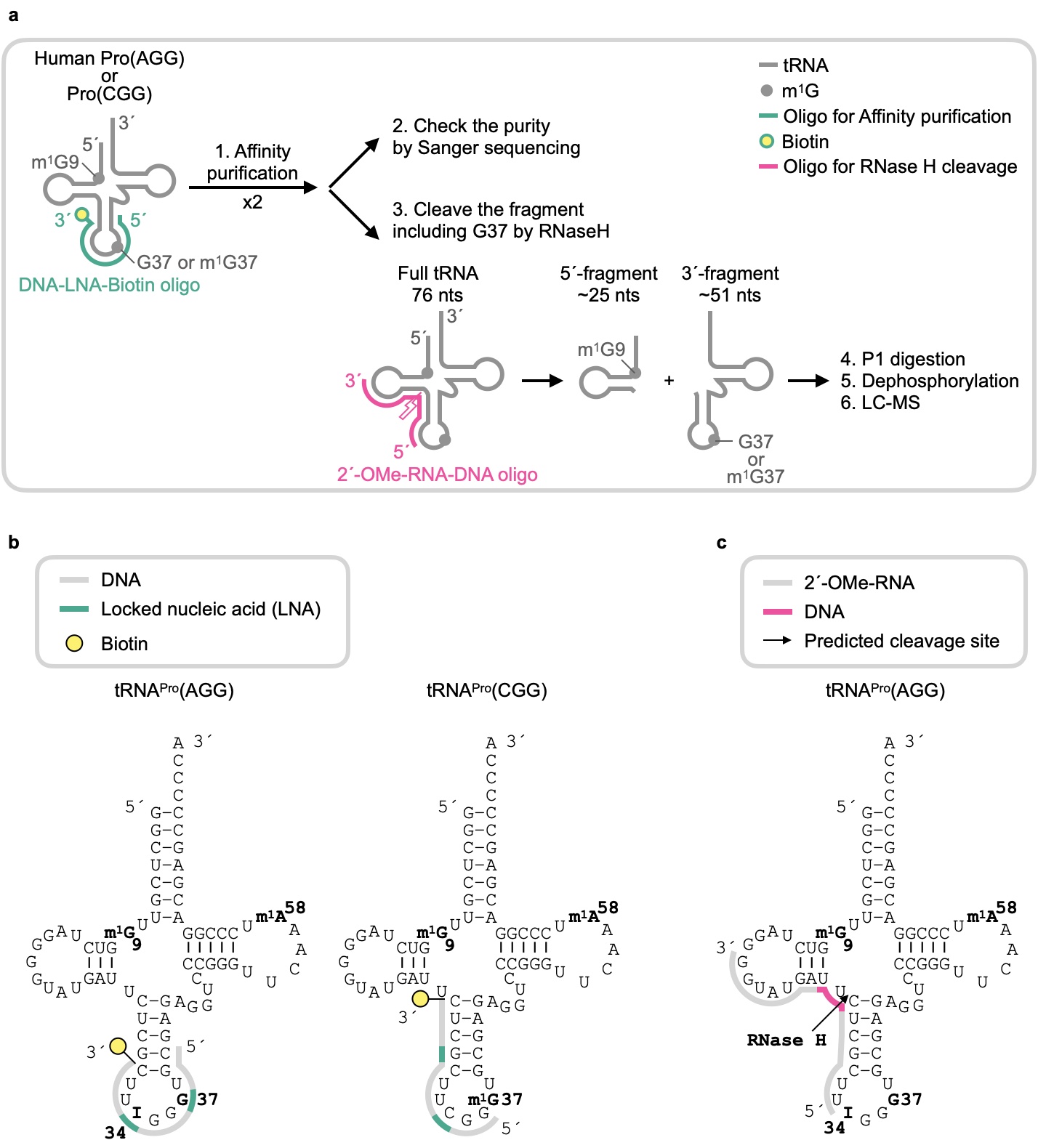

**Supplementary Figure 6.** **Sample preparation for LC-MS analysis.**

**a.** The workflow of sample preparation of human Pro(AGG) and Pro(CGG) for LC-MS analysis. Step 1: Each tRNA was purified from total RNA of K562 or yeast cells by affinity binding to a DNA-LNA-biotin oligonucleotide and by pulled-down using a streptavidin resin. This affinity purification was repeated twice. Step 2: The homogeneity of each affinity-purified tRNA was confirmed by Sanger sequencing. Step 3: Each affinity-purified tRNA was subjected to RNase H cleavage to generate a 5'-fragment of ~25 nts, which contained m^1^G9, and a 3'-fragment of ~51 nts, which contained the anticodon loop. Targeted RNase H cleavage was mediated by an oligonucleotide containing a 2'-*O*-methyl RNA/DNA backbone. Step 4: The 3'-fragment was digested to 5’-mono-phosphorylated nucleotides by P1 nuclease. Step 5: The generated 5’-mono-phosphorylated nucleotides were dephosphorylated by alkaline phosphatase. Step 5: The generated nucleosides were analyzed by LC-MS analysis. **b.** Affinity purification of human Pro(AGG) and Pro(CGG), each in the sequence and cloverleaf structure, by binding to an oligonucleotide with a DNA backbone (gray) interrupted by two LNA residues (blue). The 3’-end biotin is shown in yellow. **c.** Targeted RNase H cleavage of human Pro(AGG), shown in the sequence and cloverleaf structure, guided by an oligonucleotide consisting of both a 2’-*O*-methyl RNA backbone (gray) and a DNA backbone (red). The predicted RNase H cleavage site is shown by an arrow. The nucleotides of interest m^1^G9, I34, G37, m^1^G37, and m^1^A58 are in bold face.

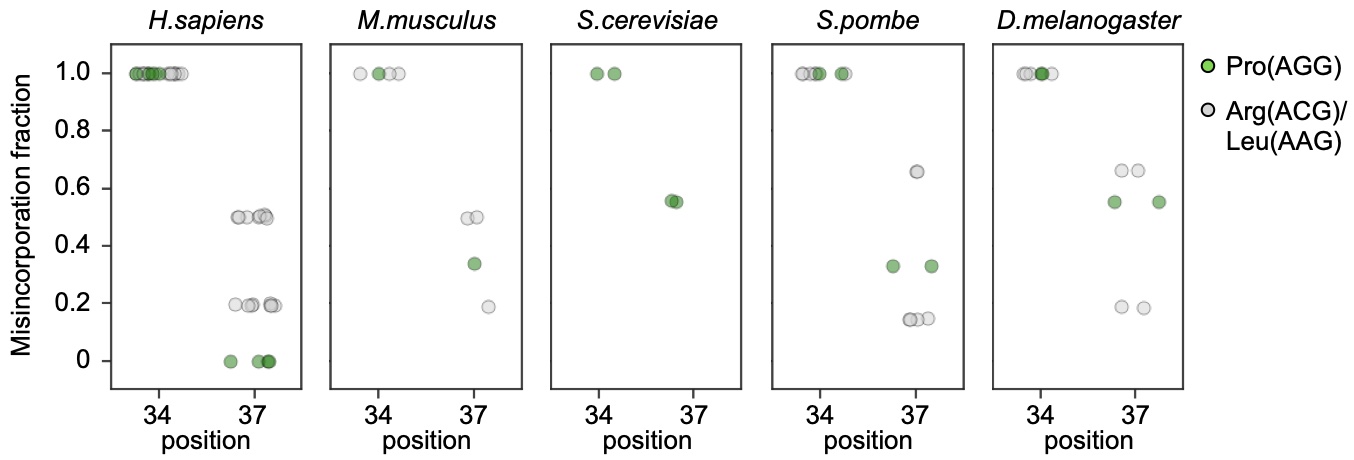

**Supplementary Figure 7. Specific** **absence of m^1^G37 from human Pro(AGG).**

Box plots indicate side-by-side direct comparison of the misincorporation frequency at position 34 and 37 in all tRNA isodecoders that contain A34 and G37 across 5 species (human *H. sapiens* (n = 4, biological replicates), mouse *M. Musculus* (n = 1), fly *D. melanogaster,* yeast *S. cerevisiae*, and yeast *S. pombe*). The data of *H. sapiens and M. musculus* are from Induro-tRNAseq, while the others are extracted from mim-tRNAseq^34^.

**Supplementary Table 1**. Sequences of DNA splint oligos used for ligation of a 3'-barcoded adaptors. Barcodes and ribonucleic acids are indicated in italic and bold.

| Barcode | 3´-RNA/DNA barcoded adaptor (5´ to 3´) | DNA splint oligo (5´ to 3´) |
| --- | --- | --- |
| 1 | phosphate-*GAU****A*TCGT**CAAGATCGGAAGAGCACACGTCTGAAC-amino | CGATCTTG**ACGAT**ATCTGGN |
| 2 | phosphate-*GAU****A*GCTA**CAAGATCGGAAGAGCACACGTCTGAAC-amino | CGATCTTG**TAGCT**ATCTGGN |
| 3 | phosphate-*GAU****G*CATA**CAAGATCGGAAGAGCACACGTCTGAAC-amino | CGATCTTG**TATGC**ATCTGGN |
| 4 | phosphate-*GAU****U*CTAG**CAAGATCGGAAGAGCACACGTCTGAAC-amino | CGATCTTG**CTAGA**ATCTGGN |

**Supplementary Table 2**. Sequences of oligos for PCR. Asterisks indicate a phosphorothioate backbone. Barcodes are indicated in bold.

|  | Sequence (5´ to 3´) |
| --- | --- |
| Multiplex primer | AATGATACGGCGACCACCGAGATCTACACTCTTTCCCTACACGACGCT*C |
| Barcoded primer I | CAAGCAGAAGACGGCATACGAGAT**ACATCG**GTGACTGGAGTTCAGACGTGT*G |
| Barcoded primer II | CAAGCAGAAGACGGCATACGAGAT**ACATCG**GTGACTGGAGTTCAGACGTGT*G |
| Barcoded primer III | CAAGCAGAAGACGGCATACGAGAT**GCCTAA**GTGACTGGAGTTCAGACGTGT*G |
| Barcoded primer IV | CAAGCAGAAGACGGCATACGAGAT**TGGTCA**GTGACTGGAGTTCAGACGTGT*G |
| Barcoded primer V | CAAGCAGAAGACGGCATACGAGAT**CACTGT**GTGACTGGAGTTCAGACGTGT*G |
| Barcoded primer VI | CAAGCAGAAGACGGCATACGAGAT**ATTGGC**GTGACTGGAGTTCAGACGTGT*G |
| Barcoded primer VII | CAAGCAGAAGACGGCATACGAGAT**TCAAGT**GTGACTGGAGTTCAGACGTGT*G |

**Supplementary Table 3**. Sequences of DNA oligos used for affinity purification and RNase H cleavage of tRNA. LNAs are in bold and indicated by “+” before the nucleotide. The 3’-biotin tag is denoted by “/BioTEG/”. Note that 2’-OMe RNAs are indicated by “m” before the nucleotide.

|  | Sequence (5´ to 3´) |
| --- | --- |
| Human Pro(AGG)/(CGG)expressed in yeast | GGCTCGTCCGGGATTTGAAC/BioTEG/ |
| Human Pro(AGG) | GCA**+C**CC**+C**AAG/BioTEG/ |
| Human Pro(CGG) | CC**+G**AAG**+C**GAG/BioTEG/ |
| RNase H for Pro(AGG)/(CGG) | mAmAmGmCmGmAGAAmUmCmAmUmAmCmC |

**Supplementary Table 4**. Sequences of primers used for reverse transcription and PCR amplification of affinity-purified tRNAs.

|  | Sequence (5´ to 3´) |
| --- | --- |
| Reverse transcription primer | GACACGGTACCACACAACTGGGGGCTCGTCCGGGATTTGAAC |
| PCR forward primer | GGCTCGTTGGTCTAGGGGTAT |
| PCR reverse primer | GACACGGTACCACACAACTGG |
